# Reconstruction of nitrogenase predecessors suggests origin from maturase-like proteins

**DOI:** 10.1101/2021.07.07.451390

**Authors:** Amanda K. Garcia, Bryan Kolaczkowski, Betül Kaçar

**Affiliations:** Department of Bacteriology, University of Wisconsin – Madison, USA; Department of Microbiology and Cell Science, University of Florida, Gainesville, Florida, USA

**Keywords:** nitrogenase, nitrogen fixation, maturase, ancestral sequence reconstruction, early life, historical contingency

## Abstract

The evolution of biological nitrogen fixation, uniquely catalyzed by nitrogenase enzymes, has been one of the most consequential biogeochemical innovations over life’s history. Though understanding the early evolution of nitrogen fixation has been a longstanding goal from molecular, biogeochemical, and planetary perspectives, its origins remain enigmatic. In this study, we reconstructed the evolutionary histories of nitrogenases, as well as homologous maturase proteins that participate in the assembly of the nitrogenase active-site cofactor but are not able to fix nitrogen. We combined phylogenetic and ancestral sequence inference with an analysis of predicted functionally divergent sites between nitrogenases and maturases to infer the nitrogen-fixing capabilities of their shared ancestors. Our results provide phylogenetic constraints to the emergence of nitrogen fixation and are consistent with a model wherein nitrogenases emerged from maturase-like predecessors. Though the precise functional role of such a predecessor protein remains speculative, our results highlight evolutionary contingency as a significant factor shaping the evolution of a biogeochemically essential enzyme.

**SIGNIFICANCE STATEMENT:** The origin of nitrogenase-catalyzed nitrogen fixation was a transformative event in life’s history, garnering long-term study from molecular, biogeochemical, and planetary perspectives. Reconstruction of ancestral nitrogenases suggests that the protein sequence space capable of yielding a nitrogen-fixing enzyme in the past was likely more constrained than previously thought. Specifically, here we show that nitrogenases likely evolved from ancestors that resemble maturases, homologs that today participate in nitrogenase cofactor assembly, contrary to the commonly accepted view that maturases evolved from a nitrogenase ancestor. We further submit that the molecular architecture that may have been required for nitrogenase origins was unlikely to have been shaped by the same environmental drivers often implicated in the evolution of nitrogen fixation. If this decoupling is found to be a recurring pattern in metabolic origins, then the presented results would undercut the common, systems-focused rationale of using ancient environmental conditions to explain the timing of critical and singular biogeochemical innovations in life’s past.

## INTRODUCTION

The modern biosphere is shaped by a variety of essential and ancient enzymes that have co-evolved with the Earth environment for billions of years. Though general mechanisms for the gain of novel enzymatic functions have been explored (Ohno 1970; Gerlt and Babbitt 2001; Copley 2015; Noda-Garcia et al. 2018; Copley 2021), the co-evolutionary steps toward the origins of many specific, key enzymes during Earth’s early history remain unresolved. An unavoidable task in addressing this challenge is constraining the ancestral functions of early-evolved enzyme families and their precursors (Benner et al. 2007, Kacar and Garcia 2019).

Biological nitrogen fixation is a notable example of a critical metabolic process with ancient and enigmatic origins. All life requires fixed, or bioavailable, nitrogen. For much of Earth history, this biologically vital element has primarily been acquired by organisms via the activities of nitrogenase metalloenzymes, early evolved and conserved catalysts that uniquely reduce dinitrogen (N_2_) to ammonia (NH_3_) (Hoffman et al. 2014; Einsle and Rees 2020). The evolution of nitrogenases has constrained the long-term productivity of the biosphere and has itself been shaped by the co-evolving biogeochemistry of Earth’s environment (Falkowski 1997; Glass et al. 2009; Stüeken et al. 2016; Luo et al. 2018; Allen et al. 2019; Mus et al. 2019; Garcia et al. 2020). Nitrogenases represent the only known biomolecular solution for the reduction of N_2_, a remarkable innovation given that the N≡N bond is one of the most inert in nature. The answer to how biology converged on this solution billions of years ago remains elusive (Boyd and Peters 2013; Mus et al. 2019; Garcia et al. 2020).

Insights into the origins of biological nitrogen fixation can be gained by reconstructing the protein ancestors of nitrogenases and their close homologs. Nitrogenases do not operate alone, but instead exist within a larger protein network required for their assembly and function. The closest homologs to nitrogenases are themselves a key player in this larger assembly network, serving as maturases (also referred to as assembly scaffold proteins) for the final steps in nitrogenase cofactor biosynthesis (Fay et al. 2016; Buren et al. 2020) (fig. 1a). These maturases are considered necessary in most nitrogenase assembly pathways to modify a complex metal cluster precursor that, when matured, serves as the nitrogenase active site for N_2_ reduction. (The only forms of nitrogenases that are confirmed to assemble without maturases are those that only incorporate iron into their active-site cofactors (Perez-Gonzalez et al. 2021)). Though not themselves known to reduce N_2_, these maturases reduce a variety of other substrates *in vitro* under highly reducing conditions, including C_2_H_2_, CO, and CN^-^, that also serve as alternative, non-physiological substrates of nitrogenases (Hu et al. 2010; Fay et al. 2016; Seefeldt et al. 2020). These findings establish maturases as catalytically similar to nitrogenases with the exception of their inability to reduce N_2_. The divergent protein features between nitrogenases and maturases must therefore account for this difference in N_2_-reduction capability.

**Fig. 1.**
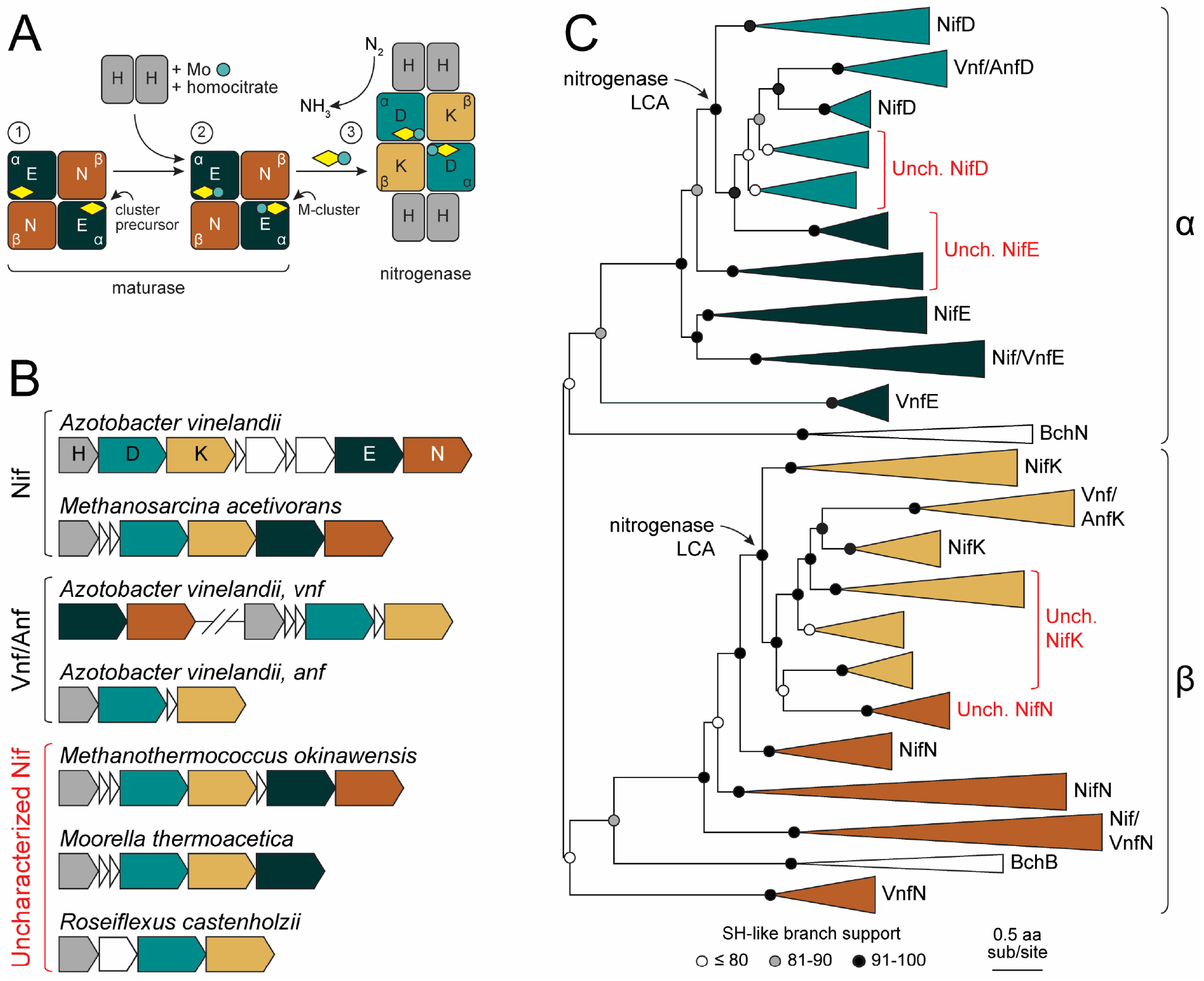
Nitrogenase and maturase functionality, genetic organization, and evolutionary history. (*A*) Simplified biosynthetic pathway for nitrogenase active-site cluster maturation and incorporation (molybdenum-dependent nitrogenase system shown). Pathway steps are indicated by circled numbers. Step 1). Maturase NifEN proteins (NifE, dark green; NifN, brown), which are α_2_β_2_ heterotetramer homologs to nitrogenase NifDK proteins (NifD, teal; NifK, yellow), are scaffolds for maturation of nitrogenase iron-sulfur cluster precursors (yellow diamond). Step 2) Cluster precursors are matured to M-clusters (yellow diamond with teal circle) by incorporation of molybdenum and homocitrate, delivered to the maturase complex by NifH proteins. Step 3) Mature M-clusters are incorporated into the nitrogenase complex, where they serve as the active sites for N_2_ reduction to NH_3_. During enzyme turnover, NifH proteins transiently interface with the nitrogenase NifDK complex to deliver electrons to the M-cluster active site. (*B*) Representative gene locus structures for molybdenum-dependent (Nif), vanadium-dependent (Vnf), and iron-dependent (Anf) nitrogenase systems, as well as for uncharacterized nitrogenase homologs (see text for discussion). Gene and intergenic region lengths are approximate. Hash marks indicate significant distance between represented genes. (*C*) Maximum likelihood phylogenetic tree (Tree-1; see text for details) built from nitrogenase and maturase protein sequences. α- or β-subunit designations for protein sequences are indicated on the right. Homologs from uncharacterized taxa are highlighted in red. Clade widths are not to scale. “LCA”: Last common ancestor.

To investigate the origins of biological nitrogen fixation, we reconstructed the evolutionary history of nitrogenases and maturases. By comparing patterns of sequence conservation between nitrogenases and maturases, we identified divergent residues that might account for their functional differentiation—namely, their ability or inability to reduce N_2_, respectively. These sequence features were then leveraged to infer the N_2_-reduction capability of reconstructed ancestral proteins and phylogenetically map the origins of biological nitrogen fixation within the nitrogenase family evolutionary history. The relative timing of the evolutionary relationship between nitrogenases and maturases is debated, with some studies suggesting that maturases are evolutionarily derived from nitrogenases (Boyd et al. 2011a). Our findings support an origin of the canonical nitrogenase clade from predecessor proteins that were unlikely to have been capable of N_2_ reduction and more closely resemble extant maturases—proteins that are today only ancillary in biological nitrogen fixation. Nitrogenases may therefore represent a case in molecular evolution where a pre-existing but already complex protein architecture, adapted to an alternative role, shaped the origins of one of the most consequential biomolecular innovations in Earth history.

## RESULTS AND DISCUSSION

### Uncharacterized homologs root canonical nitrogenases within maturase protein clades

We reconstructed the phylogenetic history of nitrogenase and maturase homologs to explore ancestral states for this protein family. There are three forms of nitrogenase—Nif, Vnf, and Anf—that each differ in the composition of their active-site iron-sulfur cluster (“M-cluster”; incorporating molybdenum, vanadium, or additional iron, respectively). The heterotetrameric (α_2_β_2_) catalytic protein component of nitrogenase (Nif/Vnf/AnfDK) has its counterpart in a homologous maturase protein complex (Nif/VnfEN; the Anf nitrogenase system does not have dedicated maturase proteins (Perez-Gonzalez et al. 2021)) (fig. 1a). We compiled a comprehensive dataset including nitrogenase Nif/Vnf/AnfDK and maturase NifEN protein sequences (fig. 1b), as well as outgroup dark-operative protochlorophyllide oxidoreductase homologs that share the α_2_β_2_ subunit arrangement (BchNB) (Fujita and Bauer 2000; Moser and Brocker 2011). Four maximum likelihood phylogenies were built to test the robustness of tree topology and downstream ancestral sequence inference to sequence sampling and alternate alignment methods (table 1): 1) 2,425 nitrogenase, maturase, and outgroup homologs aligned by MAFFT (Katoh and Standley 2013), Tree-1; 2) removal of “uncharacterized” nitrogenase and maturase homologs, Tree-2 (see definition and discussion of “uncharacterized” homologs below); 3) removal of β-subunit nitrogenase, maturase, and outgroup homologs, Tree-3; and 4) alignment with MUSCLE (Edgar 2004) instead of MAFFT, Tree-4.

**Table 1.** Nitrogenase and maturase phylogenies built in this study.

| Tree | Alignment method | Sequence Dataset |
| --- | --- | --- |
| Tree-1 | MAFFT | Nif/Vnf/AnfDK, Nif/VnfEN, BchNB |
| Tree-2 | MAFFT | Tree-1 dataset without uncharacterized homologs |
| Tree-3 | MAFFT | Tree-1 dataset without β-subunit sequences (Nif/Vnf/AnfK, Nif/VnfN, BchB) |
| Tree-4 | MUSCLE | Same as Tree-1 dataset |

General shared features across the reconstructed phylogenies include clustering of α-subunit sequences sister to β-subunit sequences (except in Tree-3, which lacks β-subunit sequences), reproducing the α_2_β_2_ structural distinction for nitrogenases and maturases (fig. 1c, supplementary fig. S1). These α- and β-subunit clades themselves each segregate into nitrogenase and maturase protein sequences. This topology is consistent with an initial gene duplication event that resulted in separate α- and β-subunits, followed by a secondary duplication event that resulted in functionally distinct nitrogenase and maturase proteins (Fani et al. 2000; Raymond et al. 2004; Boyd et al. 2011a; Boyd and Peters 2013). Within the nitrogenase clade, vanadium- and iron-nitrogenase sequences nest within molybdenum-nitrogenase clades, as has been previously observed (Raymond et al. 2004; Boyd et al. 2011a; Garcia et al. 2020). By contrast, the phylogenetic clustering of maturase sequences associated with different metal-dependent forms of nitrogenases do not reproduce this nesting pattern. Vanadium-maturase sequences are each split into two groups: one forms a small clade with a relatively long branch that diverges prior to all other nitrogenase and maturase sequences, whereas another diverges relatively recently within nitrogenase clades associated with aerobic or facultative bacteria. This topology suggests that maturases for the vanadium nitrogenase system originated independently at least twice, with one origin associated with an early divergence from ancestors of unknown function and another from a recent nitrogenase ancestor.

For the phylogenies that include them (i.e., Tree-1, Tree-3, Tree-4), several maturase-like homologs diverge prior to nitrogenases and root the latter within the maturase clade (fig. 1c, supplementary fig. S1). These homologs belong to “uncharacterized” bacterial and archaeal taxa that lack extensive experimental characterization regarding the metal dependence and N_2_- reducing capability of their nitrogenase-like proteins (McGlynn et al. 2012; Garcia et al. 2020). We obtained preliminary functional predictions for these uncharacterized, maturase-like sequences by KEGG’s BlastKOALA (Kanehisa et al. 2016), including a control subset of sequences as well that branch within canonical maturase clades. BlastKOALA returned a mix of maturase and nitrogenase annotations, even for certain control sequences (supplementary table S1). Given this discrepancy and the absence of experimental data, we assign these uncharacterized maturase-like sequences as putative maturases (rather than nitrogenases) based on three lines of evidence. First, the genes that encode these homologs in uncharacterized taxa are located closely downstream of nitrogenase-like genes, as is frequently the case with *bona fide* maturases (fig. 1b). Second, certain uncharacterized taxa (including those that are missing a NifN-like maturase subunit gene) have been shown to fix nitrogen, evidencing a functioning nitrogenase and, by extension, maturase (Mehta and Baross 2006; Dekas et al. 2009; Chen et al. 2021). Third, these homologs conserve a Cys48 residue (numbering here and hereafter from aligned *Azotobacter vinelandii* nitrogenase NifD) present in most maturase proteins (with the exception of certain VnfE homologs) and considered important for binding the cluster precursor prior to maturation (fig. 2a) (Kaiser et al. 2011). At the same time, these sequences all lack the strictly conserved nitrogenase His442 residue that ligates the active-site M-cluster and is critical for N_2_ reduction (Kim and Rees 1992; Lee et al. 1998; Li 2002; Jimenez-Vicente et al. 2018). Together, these observations suggest that the uncharacterized maturase-like homologs are unlikely to be functioning as nitrogenases and are more likely operating as canonical maturases.

**Fig. 2.**
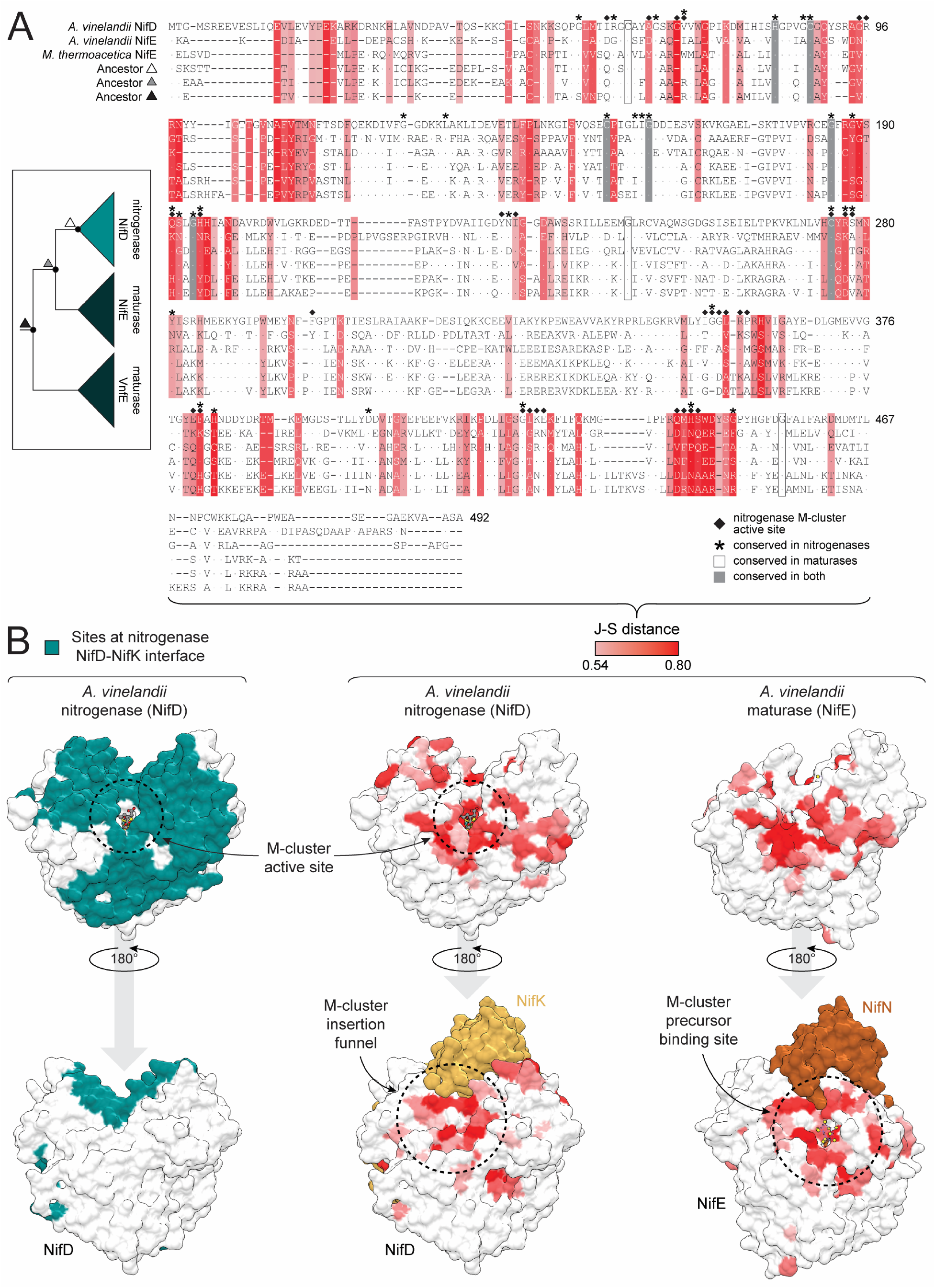
Structure and sequence maps of putative functionally divergent protein sites between nitrogenases and maturases. (*A-B*) Putative functionally divergent sites are defined as those above the 75^th^ percentile J-S distance between nitrogenase NifD and maturase NifE proteins (see text for details). The J-S distance scale applies to both (*A*) and (*B*). (*A*) Maximum likelihood ancestors (inferred from Tree-1; fig. 1c) aligned to representative extant nitrogenase (NifD) and maturase (NifE) sequences. *Moorella thermoacetica (M. thermoacetica*) NifE is an uncharacterized, putative maturase sequence. Ancestor triangle symbols correspond to labeled nodes in the simplified phylogeny (*left*) and match those in fig. 3. Nitrogenase M-cluster active-site residues are defined as those within 5 Å of the M-cluster. Dots within the alignment indicate residue identity to *Azotobacter vinelandii* (*A. vinelandii*) NifD. Site numbering based on *A. vinelandii* NifD. (*B*) Divergent sites mapped to aligned nitrogenase (*center*; *A. vinelandii* NifD, PDB 3U7Q) and maturase (*right*; *A. vinelandii* NifE, PDB 3PDI) subunit structures. Protein sites at the nitrogenase NifD-NifK interface (*left*) are defined as those within 5 Å of the NifK subunit.

Our alternate tree reconstructions demonstrate that the rooting of nitrogenases within maturase clades is primarily determined by the inclusion of uncharacterized homologs. Though the phylogenetic position of certain uncharacterized clades is ambiguous (e.g., one clade diverges immediately prior to nitrogenases in Tree-1, but prior to both nitrogenases and other maturases in Tree-3 and Tree-4), we do not observe rooting of maturases within nitrogenase sequences in any of these trees (fig. 1c, supplementary fig. S1). These topological features are also unaffected by trimming the Tree-1 alignment (supplementary fig. S2). This consistency suggests that the observed rooting pattern is robust to the exclusion of the β-subunit protein subtree (Tree-3) and variation in tested alignment methodology (Tree-4). By contrast, the exclusion of uncharacterized homologs from the sequence dataset results in reciprocal monophyly between α-subunit nitrogenase and maturase clades and nesting of β-subunit maturase sequences within nitrogenase sequences (Tree-2; supplementary fig. S1). The α-subunit topology is therefore more stable and less affected by the presence of uncharacterized homologs than the β-subunit topology, which is consistent with the comparatively low branch support values among β-subunit sequences in Tree-2.

The rooting of nitrogenases within the broader maturase protein clade might parsimoniously suggest that the functionality of the common ancestor of both protein groups more likely resembled that of extant maturases. These phylogenetic observations present a hypothesis that can be tested by evaluating sequence features of reconstructed ancestral proteins inferred to contribute to the functional divergence between nitrogenases and maturases.

### Phylogenetically divergent protein sites between extant nitrogenases and maturases map to functionally important structural regions

We performed a comprehensive, comparative sequence-structure analysis to identify protein sites that likely contribute to the functional divergence (i.e., (in)ability to reduce N_2_) between nitrogenases and maturases. Our goal was to subsequently leverage this analysis for identification of similar sites in reconstructed protein ancestors and phenotypic inference. We limited our analysis to α-subunit nitrogenase (NifD) and maturase (NifE) sequences in part due to the greater topological uncertainty within β-subunit subtrees across our phylogenetic reconstructions (fig. 1c, supplementary fig. S1). In addition, α-subunit sequences host the active-site M-cluster or cluster precursor. We therefore expected that functional differences between nitrogenases and maturases are more likely to be modulated by sequence- and structural-level differences between α-subunit proteins.

To predict functionally divergent protein sites, we first calculated the amino acid frequency distributions of each alignment column for nitrogenase and maturase sequences (Materials and Methods). The Jensen-Shannon (J-S) distance between the two protein groups for every alignment column was calculated, where larger distances indicate greater divergence for that site between nitrogenases and maturases. We estimated the expected distribution of site-wise J-S distances by randomly partitioning our protein sequences into 2 groups 10,000 times and calculating site-wise J-S distances from each random partition. The p-value for each site’s J-S distance from the nitrogenase-maturase partition was calculated from the distribution of J-S distances for that site across random sequence partitions. We defined functionally divergent sites as those exceeding the 75^th^-percentile distance across all alignment columns, as well as having an FDR-corrected *p*-value <0.0001. This analysis was repeated for all four alignments used to build phylogenetic trees (table 1), yielding 117 (116 for the Tree-4 alignment) predicted sites (supplementary table S2).

Putative functionally divergent sites identified from the sequence alignments cluster with known functionally important structural regions of the nitrogenase subunit (fig. 2b). These include the M-cluster active site for N_2_ reduction, the interface between nitrogenase NifD and NifK subunits, and the M-cluster insertion funnel (Hu et al. 2008). This observed correlation suggests that the set of putative functionally divergent sites is likely enriched for positions contributing to the functional divergence between nitrogenases and maturases. Six sites are of the most divergent across all sequence alignments: 69, 189, 362, 383, 440, and 444 (supplementary table S2). Two additional sites, 442 and 445, are highly divergent in alignments for Tree-1, Tree-2, and Tree-3, but not for Tree-4, likely because Tree-4 was constructed by a different alignment method that would have impacted downstream distance calculations (table 1). All are in or proximal to the nitrogenase active site or M-cluster insertion funnel. Some have specific inferred or experimentally determined functional roles in nitrogenases, including as an M-cluster ligand (site 442) (Kim and Rees 1992), a “lock” to hold the M-cluster within the active site (site 444) (Hu et al. 2008; Solomon et al. 2020), and a “lid” at the cluster insertion funnel opening (site 362) (Hu et al. 2008). Finally, our set of divergent sites includes the maturase cluster precursor ligand, site 48 (Kaiser et al. 2011). The clustering of highly divergent sites with these protein regions highlights the differential interaction of nitrogenases or maturases with the M-cluster. Specifically, maturase function does not require the conservation of residues that permit the insertion and stabilization of the cluster within the active site (Hu et al. 2008), nor the fine-tuning of residues in the active site for catalysis.

In addition to divergent sites, we identified several conserved residues among nitrogenase and maturase sequences (fig. 2a). His83, Cys88, Cys154, Gly160, Gly185, Gly194, and Cys275 residues are conserved in both nitrogenases and maturases. These sites are likely essential to both groups and arose prior to their evolutionary divergence. 23 sites are conserved only in nitrogenases, compared to just three sites that are conserved only in maturases: Cys62, Gly246, and Gly455. However, these three residues are still present in most nitrogenase sequences. The greater number of uniquely conserved residues in nitrogenases relative to maturases may reflect the stronger selective constraint associated with N_2_-reduction functionality.

### Putative functionally divergent sites of oldest ancestors resemble extant maturases more than nitrogenases

With a set of predicted functionally divergent protein sites between extant α-subunit nitrogenase and maturase proteins, we used a probabilistic approach to compare features of ancestral proteins inferred from all reconstructed phylogenies (table 1, supplementary fig. S1). For divergent sites, J-S distances were calculated between ancestral amino acid posterior probability distributions and extant amino acid frequency distributions for either extant nitrogenase or maturase homologs. These distance scores were then normalized to yield a value between -1 and +1, here called the “D-score” (supplementary table S3). Positive D-scores indicate greater similarity to α-subunit nitrogenase NifD homologs and negative scores indicate greater similarity to NifE homologs (see Materials and Methods for additional details). D-scores were averaged across divergent sites for each ancestral node.

Ancestral protein sequences inferred from oldest phylogenetic nodes on average resemble extant maturase homologs more than nitrogenase homologs at predicted functionally divergent sites. Mean D-scores across divergent sites for nodes ancestral to all nitrogenase and maturase homologs (fig. 3, “nitrogenase/maturase last common ancestor 1”) range between -0.08 and 0.00. These values are low-magnitude relative to the full range of mean D-scores across all ancestral nodes (∼0.2 to +0.2), which is expected due to the mixing of both nitrogenase- and maturase-like sequence features at oldest nodes. Nevertheless, the primarily negative mean D-score values at these nodes indicate greater sequence-level similarity to maturase Nif/VnfE sequences than to nitrogenase Nif/Vnf/AnfD sequences. The ambiguity of the Tree-3 node (D-score = 0.00) may result from the removal of β-subunit sequences for this phylogeny that would otherwise form an outgroup to the α-subunit clade and constrain ancestral sequence composition at this node. At more recent ancestral nodes that exclude early diverged VnfE clades (which have long, less highly supported branches and, thus, ambiguous evolutionary context; “nitrogenase/maturase last common ancestor 2”), mean D-scores range between -0.10 and +0.02. The only tree that yields positive D-scores for these more recent nodes is that reconstructed from the MUSCLE alignment (Tree-4), indicating that this analysis is more sensitive to the tested alignment method than sequence sampling. The sensitivity of ancestral sequence inference to alignment method has been observed previously with simulated data, also finding that MUSCLE produces less accurate inferences than MAFFT (Vialle et al. 2018). Finally, nodes associated with the “nitrogenase last common ancestor” yield positive D-scores for all tree, and therefore resemble nitrogenases more than maturases at predicted functionally divergent sites.

**Fig. 3.**
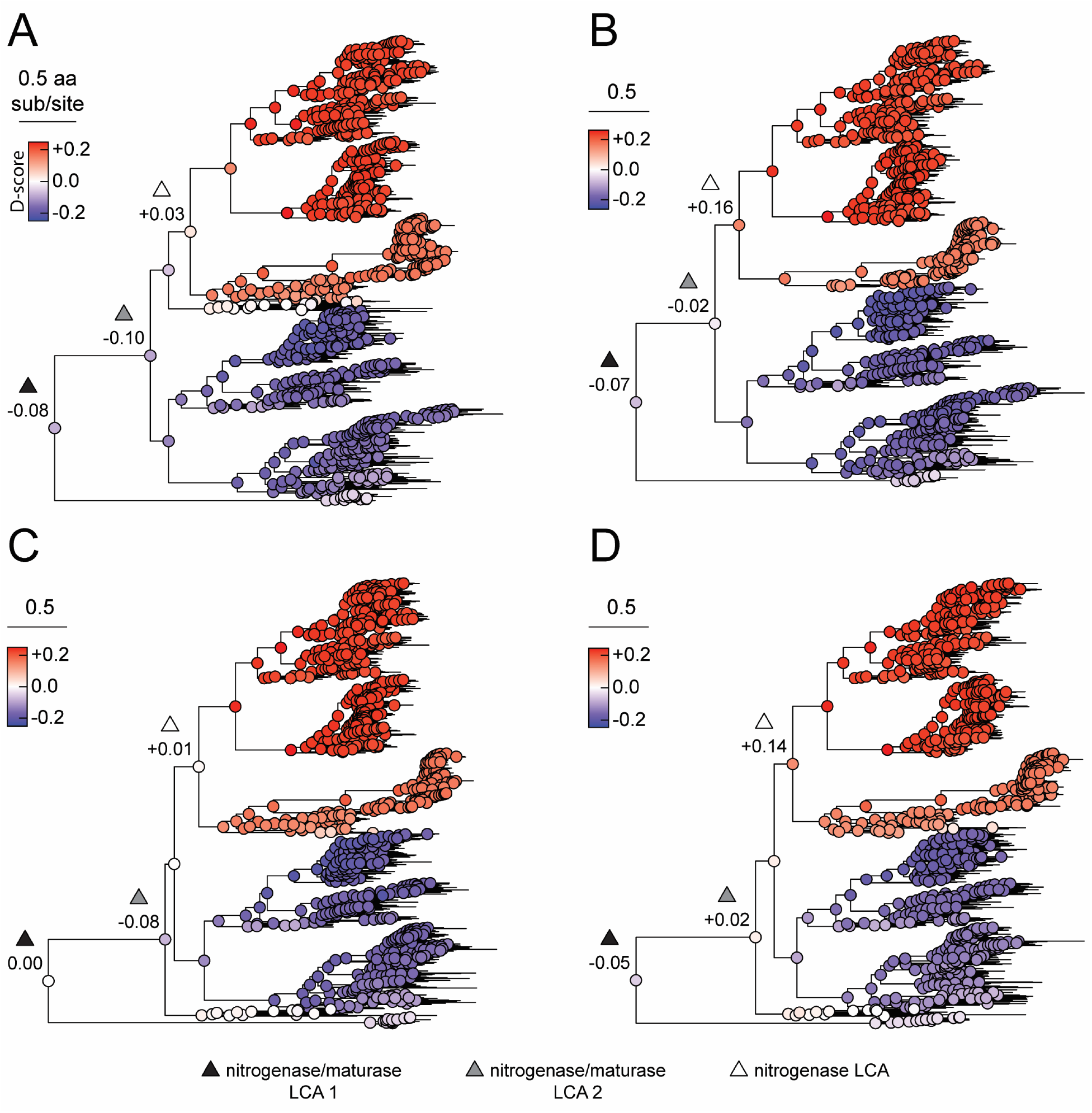
Ancestral sequence similarity to extant nitrogenases or maturases, mapped across four alternate phylogenies: (*A*) Tree-1, (*B*) Tree-2, (*C*) Tree-3, and (*D*) Tree-4. Similarity is expressed as the “D-score” parameter, where a positive D-score (red) indicates greater similarity to extant nitrogenases, and a negative D-score (blue) indicates greater similarity to extant maturases (see Materials and Methods). D-scores were averaged across putative functionally divergent sites for each ancestral node. Nodes are labeled nitrogenase/maturase last common ancestor (“LCA”) 1— including early diverged VnfE homologs (black triangle; see text for discussion); nitrogenase/maturase LCA 2—excluding early diverged VnfE homologs (grey triangle); and nitrogenase LCA (white triangle), along with their mean D-scores.

In addition to surveying mean ancestral similarity to extant nitrogenases or maturases across all putative functionally divergent sites, we also investigated site-wise D-score values, some of which have been experimentally determined to contribute to either nitrogenase or maturase function. These site-wise D-scores were assessed specifically along the phylogenetic transect between the last common ancestor of nitrogenases and the last common ancestor of all nitrogenase and maturases (supplementary table S3). Certain divergent sites that become “nitrogenase-like” early (i.e., prior to the nitrogenase ancestor) include site 195 (important for N_2_ substrate binding (Kim et al. 1995)), site 444 (locks the M-cluster in the nitrogenase active site (Hu et al. 2008; Solomon et al. 2020)), and site 359 (helps form the cluster insertion funnel (Hu et al. 2006)). By contrast, site 48 (involved with L-cluster binding in maturases (Kaiser et al. 2011)), as well as sites 361 and 362 (involved with M-cluster insertion at the nitrogenase active site (Hu et al. 2008)), remain primarily “maturase-like” until the nitrogenase last common ancestor.

The site 442 histidine M-cluster ligand conserved across all extant canonical nitrogenase proteins is considered critical for nitrogenase activity (Kim and Rees 1992; Lee et al. 1998; Li 2002; Jimenez-Vicente et al. 2018) and, notably, remains maturase-like prior to the nitrogenase last common ancestor (mean D-score ≈ -0.40 across alternate alignments, compared to a minimum D-score of -0.76 for all divergent sites; supplementary table S3). Site 442 is only not predicted to be functionally divergent for the tree reconstructed by a MUSCLE alignment (Tree-4). This difference is likely due to an inferred homology, unique to the MUSCLE alignment, between the nitrogenase His442 residue and a frequently observed histidine residue in maturases. However, the other alignments, which instead infer homology with a neighboring arginine residue in many maturases (fig. 2a), is supported by studies which indicate that this arginine residue is structurally aligned to nitrogenase His442 (Kaiser et al. 2011). Thus, it is likely that the MUSCLE alignment is erroneous at this site, consistent with the observed reduced accuracy of MUSCLE compared to MAFFT for ancestral sequence inference (Vialle et al. 2018). Due to its functional significance and conservation among extant nitrogenases, the appearance of a histidine residue at site 442 may have been critical for the origins of N_2_ reduction.

### Nitrogenases likely originated from a non-N_2_-reducing maturase-like protein with possible biosynthetic or alternate catalytic roles

Our exploration of nitrogenase and maturase ancestry, coupled with the investigation of global sequence and structural features that account for their functional divergence, is generally consistent with the hypothesis that the last common ancestor of nitrogenase and maturase proteins was unlikely to have functioned as a nitrogenase. This inference is also supported by the absence of residue-level similarity at divergent sites that have empirically been shown to be critical for nitrogenase function. These results are robust to phylogenetic uncertainty stemming fromsequence sampling of early diverged uncharacterized lineages and incorporates statistical uncertainty associated with ancestral sequence inference. Though we find that an alternative alignment method, MUSCLE, does modulate these sequence-based functional inferences at one ancestral node (fig. 3), there is reason to suspect reduced alignment accuracy given probable misalignment of at least one key nitrogenase residue and decreased performance in simulated data (Vialle et al. 2018) (see above).

Conservatively, our results suggest that phylogenetic inference of ancient nitrogen fixation can reliably extend only to within the canonical nitrogenase clade, and that inferences of earlier nitrogen fixation activity require further evidence. More definitive assessment of the N_2_-reduction capability of the common ancestor of nitrogenases and maturases awaits experimental investigation, which can be directly achieved through the laboratory resurrection of ancestral proteins inferred in this study (Thornton 2004; Benner 2017; Garcia and Kacar 2019). Nevertheless, a model that posits a maturase-like ancestry for nitrogenases deviates from existing hypotheses regarding their early evolution (fig. 4). Previous models are based on parsimonious interpretations of nitrogenase and maturase phylogenetic topology that is not observed in the trees reconstructed here. For instance, previous studies root the maturase clade within nitrogenase sequences, suggesting that the former evolved via gene duplication of nitrogenase ancestors (Boyd et al. 2011a; Boyd and Peters 2013). An updated phylogenetic analysis incorporating the Boyd et al. (2011a) dataset, lacking uncharacterized sequences, produces a topology similar to Tree-2 where nitrogenase and maturase α-subunit clades are reciprocally monophyletic (supplementary fig. S2). However, with the inclusion of uncharacterized nitrogenase and maturase homologs, maturases instead root nitrogenases, and support a model wherein nitrogen fixation is instead a derived feature of a maturase-like ancestor. Our reconstruction of ancestral states within the nitrogenase and maturase phylogeny provides additional constraints on ancestral phenotypes within a maximum likelihood framework, extending beyond inferences drawn from phylogenetic topology alone. These results constrain the likely origin of nitrogen fixation to a relatively well-resolved lineage within the nitrogenase/maturase topology, rather than to a deeper history that bridges nitrogenases with more distantly related homologs (e.g., coenzyme F430 biosynthesis proteins (Mus et al. 2019)).

**Fig. 4.**
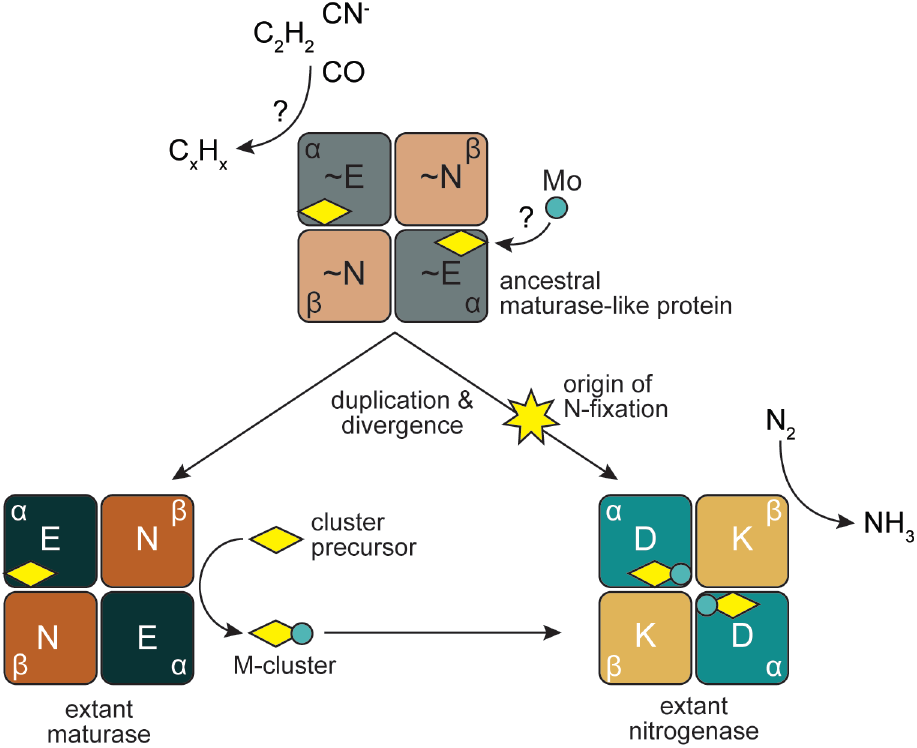
Proposed model for the origins and functional divergence of maturase and nitrogenase proteins. An ancestral maturase-like protein (∼NifEN, grey and light brown), incapable of reducing N_2_, may have otherwise reduced various carbon-containing substrates and/or played a role in cluster (yellow diamond) biosynthesis. The ancestor may have been capable of incorporating molybdenum (teal circle) into the cluster. Duplication of the encoding ancestral genes and functional divergence would then have yielded canonical maturase (NifEN, dark green and brown) and nitrogenase (NifDK, teal and yellow) proteins. Maturases would have specialized to provide a scaffold for the maturation of the nitrogenase cluster precursor (yellow diamond) to the nitrogenase active-site M-cluster (yellow diamond with teal circle). In parallel, tuning of the ancestral peptide environment along a divergent lineage would have spurred the origin of N_2_ reduction and specialization of nitrogenases for a solely catalytic role in nitrogen fixation. Protein components of the α_2_β_2_ heterotetrameric nitrogenase and maturase structures are labeled.

Our results suggest that ancestral maturase-like proteins may have provided the molecular architecture for the origin of nitrogen fixation. However, the precise functional role of such an ancestor is not clear. Candidate phenotypic attributes, shared between extant nitrogenases and maturases, may shed light on how biological N_2_ reduction evolved. For example, both are the only proteins known to bind the nitrogenase active-site M-cluster (Fay et al. 2016). In addition, both extant nitrogenases and maturases can reduce several non-physiological, alternative substrates including C_2_H_2_, CO, and CN^-^, albeit in highly reducing experimental conditions (Fay et al. 2016; Seefeldt et al. 2020). It has previously been argued that the ability of nitrogenases to reduce alternative substrates may simply be the byproduct of overcoming the significant activation barrier required for N_2_ reduction (Boyd and Peters 2013). However, combined with the evidence presented here for a maturase-like ancestry, a plausible scenario is an ancestral protein capable of reducing the shared substrates of extant nitrogenases and maturases at an M-cluster-like active site (perhaps in a role as detoxyases, as has previously been proposed ((Silver and Postgate 1973); fig. 4). This scenario would provide a stepwise path for the evolution of nitrogenases from ancestral proteins capable of catalyzing less ATP-intensive reactions (Hu et al. 2010), and requires only residue-level tuning of an already complex peptide environment to achieve the earliest whiffs of N_2_ reduction. Our reconstruction of ancestral residues at predicted functionally divergent sites provides possible mutational trajectories toward the evolution of nitrogenases.

The origin of nitrogenases from proteins involved with cofactor biosynthesis such as the maturases studied here might be expected given the prevalence of biosynthetic proteins associated with the broader family of nitrogenase-like homologs. These include chlorophyll biosynthesis proteins used as phylogenetic outgroups in our study (Fujita and Bauer 2000; Moser and Brocker 2011), as well as coenzyme F430 biosynthesis proteins that are conserved among methanogens (Staples et al. 2007; Zheng et al. 2016). In addition, more distantly related and poorly studied nitrogenase-like homologs may have roles in assembly of hydrogenase metalloclusters and metal transport (reviewed in Ghebreamlak and Mansoorabadi (2020)). It is not clear whether a hypothetical, maturase-like nitrogenase ancestor may have had a central function in cofactor biosynthesis as its descendants and several related homologs do today. Other distantly related nitrogenase-like homologs have putative catalytic roles, including a recently reported homolog suggested to participate in a methionine salvage pathway that forms ethylene (North et al. 2020). It is possible that a maturase-like predecessor would have been promiscuous, a suggested general feature of early-evolved proteins (Copley 2015; De Tarafder et al. 2021), and capable of both providing a scaffold for cluster maturation as well as catalysis using the same matured cluster. Gene duplication and divergence might then have subsequently specialized maturases to only function as a scaffold and evolve residues that permit the release of the matured cluster. In parallel, nitrogenases would have specialized to function in a solely catalytic role.

The possible capability of an ancestral maturase-like protein to bind an M-cluster-like cofactor would have important implications for the coevolutionary trajectory of nitrogenases and the biogeochemical environment (Anbar and Knoll 2002; Glass et al. 2009; Boyd et al. 2011b). A defining role of extant maturases in the molybdenum-dependent nitrogenase system is to provide a scaffold for the incorporation of molybdenum into the active-site M-cluster. Thus, the origin of maturase proteins has previously been suggested to coincide with the origin of molybdenum dependence in nitrogenases (Boyd et al. 2011a). However, if maturase-like proteins predate nitrogenases, the molybdenum-containing M-cluster itself may predate nitrogenases as well (fig. 4). Inferences for the age of nitrogenases extend to more than 3 billion years ago (Stueken et al. 2015; Parsons et al. 2020) when molybdenum in Earth’s oceans was likely exceedingly scarce (Anbar 2008). A molybdenum-incorporating maturase-like protein existing prior to 3 billion years ago would suggest that the bulk geochemistry of the early Earth environment may not necessarily have provided strict constraints on enzyme evolution (Garcia et al. 2020), and would be in agreement with geochemical evidence of early molybdenum-dependent nitrogenase activity (Stueken et al. 2015). Rather, localized environments may have provided sufficient molybdenum for the function of nitrogenases and their predecessors (Stueken et al. 2015), or molybdenum may have simply been selected despite its scarcity due to its invaluable chemical features.

Another possibility is that earliest maturases-like ancestors did not incorporate molybdenum, but rather the iron-only cluster precursor that is matured to the M-cluster today (Mus et al. 2019).

### The role of contingency and subsumed complexity in nitrogenase evolution

The alternative model of nitrogenase origins that we propose exemplifies a case for molecular evolution in which a novel and consequential metabolic function was built off a pre-existing, complex molecular architecture. An open question is whether the pre-existing complexity and functional role of a maturase-like protein was needed to evolve an enzyme capable of reducing N_2_. The role of evolutionary contingency in shaping biological diversity has long been examined (Gould 1989; Vermeij 2006; Blount et al. 2018), particularly to envision future evolutionary scenarios or alternate trajectories on other worlds characterized by distinct environmental parameters (Kaçar and Gaucher 2012). Regarding the evolution of biological nitrogen fixation in particular, it has been argued that necessity—i.e., the need for bioavailable sources of nitrogen— or environmental geochemistry likely controlled the timing of early nitrogenase evolution and diversification on Earth (Navarro-Gonzalez et al. 2001; Anbar and Knoll 2002; Mus et al. 2018). However, another possibility is that the origin of biological nitrogen fixation required the subsumed complexity of a protein predecessor (Adam et al. 2018), which was initially positively selected for an entirely different metabolic role. The origin of nitrogenases thus may not have occurred (or may have been significantly delayed) without a suitable protein on which to build, despite the scarcity of bioavailable nitrogen, an abundance of possible metal cofactors, or both. Testing these possibilities would require experimentally replaying the evolutionary path that led to the origin of biological nitrogen fixation.

There is no evidence that Terran biology evolved nitrogen fixation more than once. Whether this is the product of very exceptional circumstances of origination or of a survivorship bias so pronounced that there is only one remaining functional example supported across all of biology, nitrogenases are therefore on par with other singular molecular-level innovations such as the ribosome (Fox 2010) and oxygenic photosynthesis (Blankenship and Hartman 1998). The existing variation among different metal-dependent forms of nitrogenase enzymes do not constitute truly independent evolutionary experiments, but variations on a theme that was determined and uniquely constrained by the common ancestral form. Even within this narrow range of constraints, the degree to which contingent amino-acid substitutions shaped the diversification of nitrogenase metal preference and specificity remains unknown.

Identifying the singular circumstances that left Earth with a single common ancestor for all nitrogenase function may be critical for understanding the pervasiveness of nitrogen fixation as a universal biological capability. This is particularly important for assessing whether metal availability significantly guides protein evolution or whether other internal biophysical constraints lead to background-dependent, epistatic interactions (Williams and da Silva 1996; Anbar and Knoll 2002; Moore et al. 2017; Smethurst and Shcherbik 2021). Efforts to generate artificial nitrogenases and nitrogenase metalloclusters (Tanifuji et al. 2015; Sickerman et al. 2017) may expand the suite of molecular structures capable of reducing N_2_, but biotic experiments integrating gene regulatory and protein-protein interaction constraints are needed to test different macroevolutionary hypotheses of nitrogenase emergence. A survey of such functional constraints on nitrogenase and maturase predecessors could reveal the sequence of biomolecular functions conducive for the evolution of nitrogen fixation, which could then be integrated into a more comprehensive accounting of internal selective forces, geochemical features and planetary environments that can host similar evolutionary pathways (Kacar et al. 2020).

Perhaps most intriguingly, our results suggest that nitrogen fixation may have emerged from natural selection acting on a maturase-like protein whose ancestral function was largely decoupled from extracellular conditions frequently implicated as drivers for the origins of nitrogen fixation. This scenario would thus cast the origin of nitrogen fixation as an act of extreme contingency bordering on happenstance, betraying its utility as one of the most evolutionarily significant and biologically limiting metabolic pathways on Earth. If borne out by further study or found to be a recurring pattern for other critical molecular innovations, enzyme origins largely decoupled from putative environmental drivers may severely compromise the soundness of systems-focused hypotheses that tie organismal or ecological need to bulk geochemical substrate or cofactor availabilities. The paleobiology of molecular innovations would require disciplinary approaches and conceptual foundations quite distinct from the study of their more recent evolution.

## MATERIALS AND METHODS

### Phylogenetic reconstruction and ancestral sequence inference

Protein homologs were identified from the National Center for Biotechnology Information non-redundant protein database by BLASTp (Camacho et al. 2009) with an expect value cutoff of <1e- 5 (accessed January 2020). Query sequences from *Azotobacter vinelandii* (NifD: WP_01270336, NifK: WP_012698833; NifE: WP_012698838, NifN: WP_012698839) were used for nitrogenase and maturase homolog identification, and sequences from *Synechococcus elongatus* (BchN: WP_126148028, BchB: WP_126147769) for outgroup dark-operative protochlorophyllide oxidoreductase homolog identification. Sequences from this relatively permissive BLASTp search were aligned using MAFFT v7.450 (Katoh and Standley 2013) to build a preliminary phylogeny with FastTree v2.1.11 (Price et al. 2010). Putative nitrogenase homologs were identified based on previously published phylogenies (Boyd et al. 2011a; Garcia et al. 2020) as well as sequence features known to be critical for N_2_ reduction (e.g., Cys275, His442). Putative maturase homologs were only retained if the encoding genes were co-localized with nitrogenase genes in the same genome. Finally, sequences in overrepresented clades were pruned to obtain a roughly equal number of nitrogenase and maturase versus outgroup sequences, so as not to bias subsequent ancestral sequence inference.

A final untrimmed MAFFT alignment was used as input for phylogenetic reconstruction by RAxML v8.2.11 (Stamatakis 2014) using 100 rapid bootstrap searches and the best-fit LG+G+F evolutionary model determined by ModelFinder (Kalyaanamoorthy et al. 2017) (an additional phylogeny was also built from an alignment trimmed by TrimAl (Capella-Gutierrez et al. 2009) (supplementary fig. S2)). The tree was further optimized by applying nearest-neighbor-interchanges before calculation of SH-like branch support values (Guindon et al. 2010), resulting in the final phylogeny, Tree-1. Additional trees incorporated in ancestral sequenc inference were generated by altering sequence sampling or alignment with MUSCLE v3.8.425 (Edgar 2004) instead of MAFFT (see Results and Discussion, table 1). Finally, the Bayesian phylogenetic analysis by Boyd et al. (Boyd et al. 2011a) was replicated using their reported methods (supplementary fig. S2).

Ancestral sequences were inferred by maximum likelihood marginal reconstruction in PAML v4.9j (Yang 2007) using the same evolutionary model parameters described above for RAxML. Sequence gaps were reconstructed in PAML using the binary encoding approach described in Aadland et al. (2019). Briefly, the protein sequence alignment was recoded as a ‘presence-absence’ alignment matrix, and the posterior-probability of the presence (amino-acid residue) or absence (gap) state at each position in each ancestral sequence was calculated using maximum-likelihood reconstruction, assuming a binary character model with state frequencies inferred by maximum likelihood. All phylogenetic data, including sequence alignments, trees, and ancestral sequence inference outputs can be found at https://github.com/kacarlab/maturase2021.

### Prediction of functionally divergent protein sites between nitrogenases and maturases

For each position in the sequence alignment, we calculated the Jensen-Shannon (J-S) distance between the amino-acid frequency distribution estimated from extant nitrogenase Nif/Vnf/AnfD sequences in the alignment and that estimated from extant maturase Nif/VnfE sequences, with nitrogenase and maturase sequences being defined based on monophyly in the tree topology and gene location within the *nif*, *vnf*, or *anf* loci (see Results and Discussion, fig. 1b). Briefly, the J-S distance is calculated as the average Kullback-Leibler divergence, or “relative entropy,” which estimates the loss of information when one frequency distribution is used to represent another. Intuitively, the site-wise J-S distance between nitrogenase and maturase amino-acid frequency distributions describes how dissimilar the distribution of extant amino-acids is between nitrogenase and maturase sequences for each alignment site.

We estimated the expected distribution of site-wise J-S distances, given our sequence alignment, by randomly partitioning the alignment into two sequence groups of sizes equivalent to the sizes of our actual nitrogenase and maturase groups, respectively, and calculating site-wise J-S distances between these randomly partitioned groups. We performed 10,000 random partitions and site-wise J-S distance calculations. For each alignment column *i*, we calculated the probability of observing J-S(nitrogenase, maturase)*_i_*, given the distribution of J-S-distances at column *i* in the randomly partitioned dataset (i.e., *p*-value). We enriched for highly divergent positions using an FDR-corrected *p*-value cutoff of 0.0001. We additionally excluded any sites with J-S distances in the lower 75^th^ percentile of the J-S distance distribution across all alignment positions.

### Probabilistic assessment of nitrogenase-like ancestral sequence features

Nitrogenase-like ancestral sequence features were assessed by incorporating the statistical uncertainty of ancestral sequence inference into comparisons between those of extant nitrogenases and maturases. For putative functionally divergent protein sites identified as described above, J-S distances were calculated between the ancestral amino acid posterior probability distributions and the extant amino acid frequency distributions across either extant nitrogenase (Nif/Vnf/AnfD) or maturase (Nif/VnfE) clades. These distance values were then normalized to yield a value between -1 and +1 indicating the relative similarity of an ancestral protein site to a homologous extant nitrogenase site, here called the “D-score” (i.e., similarity to the nitrogenase D-subunit):

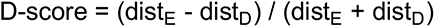

where dist_D_ is the J-S distance between ancestral and maturase E-subunit sites, and dist_E_ is the J-S distance between ancestral and nitrogenase D-subunit sites. D-scores were analyzed on a site-wise basis as well as averaged across the length of each ancestral sequence for all constructed phylogenies. All data and scripts related to the prediction of functionally divergent sites prediction and D-score calculations can be found at https://github.com/kacarlab/maturase2021.

## ACKNOWLEDGEMENTS

This work was supported by the National Aeronautics and Space Administration Postdoctoral Program to A.K.G., the National Science Foundation Division of Molecular and Cellular Biosciences (1817942 to B. Kolaczkowski), the National Science Foundation Emerging Frontiers Program (1724090 to B. Kaçar), the National Aeronautics and Space Administration Early Career Faculty Award (80NSSC19K1617 to B. Kaçar), and the Metal Utilization and Selection across Eons (MUSE) Interdisciplinary Consortium for Astrobiology Research, sponsored by the National Aeronautics and Space Administration Science Mission Directorate (19-ICAR19_2-0007 to B. Kaçar). We thank Eric Boyd and Jennifer Glass for the helpful discussions.

## DATA AVAILABILITY STATEMENT

All phylogenetic datasets and scripts are available at https://github.com/kacarlab/maturase2021.

