## Supplementary Information for "Reconstruction of nitrogenase predecessors suggests origin from maturase-like proteins"

**Table S1.** BlastKOALA (Kanehisa et al. 2016) functional predictions of uncharacterized NifE- and NifN-like homologs. Blank cells indicate that no annotation was found. KO: KEGG Orthology identifier.

| Type | Query | KO | Definition | Score | Second best KO | Second best score |
| --- | --- | --- | --- | --- | --- | --- |
| NifE-like | Ammonifex sp. | K02587 | nifE; nitrogenase molybdenum-cofactor synthesis protein NifE | 116 |  |  |
| NifE-like | Calderihabitans maritimus | K02587 | nifE; nitrogenase molybdenum-cofactor synthesis protein NifE | 135 |  |  |
| NifE-like | Caldicellulosiruptor saccharolyticus | K02587 | nifE; nitrogenase molybdenum-cofactor synthesis protein NifE | 108 | K02586 | 83 |
| NifE-like | Candidatus Desulforudis audaxviator MP104C |  |  |  |  |  |
| NifE-like | Clostridia bacterium |  |  |  |  |  |
| NifE-like | Desulfobacteraceae bacterium Eth-SRB1 | K02587 | nifE; nitrogenase molybdenum-cofactor synthesis protein NifE | 85 | K02586 | 53 |
| NifE-like | Desulfoglaeba alkanexedens |  |  |  |  |  |
| NifE-like | Methanocaldococcus infernus | K02587 | nifE; nitrogenase molybdenum-cofactor synthesis protein NifE | 218 |  |  |
| NifE-like | Methanococcus aeolicus | K02586 |  | 231 |  |  |
| NifE-like | Methanofervidicoccus sp. A16 | K02587 | nifE; nitrogenase molybdenum-cofactor synthesis protein NifE | 230 | K02587 | 19 |
| NifE-like | Methanothermococcus okinawensis | K02586 |  | 306 |  |  |
| NifE-like | Methanothermococcus thermolithotrophicus | K02586 |  | 326 |  |  |
| NifE-like | Methanotorris igneus | K02587 | nifE; nitrogenase molybdenum-cofactor synthesis protein NifE | 205 |  |  |
| NifE-like | Moorella thermoacetica | K02587 | nifE; nitrogenase molybdenum-cofactor synthesis protein NifE | 166 |  |  |
| NifE-like | Syntrophothermus lipocalidus | K02587 | nifE; nitrogenase molybdenum-cofactor synthesis protein NifE | 201 | K02586 | 29 |
| NifE-like | Thermacetogenium phaeum | K02587 | nifE; nitrogenase molybdenum-cofactor synthesis protein NifE | 223 |  |  |
| NifE-like | Thermoanaerobacterium sp. RBIITD | K02587 |  |  |  |  |
| NifE-like | Thermodesulfitimonas autotrophica | K02587 | nifE; nitrogenase molybdenum-cofactor synthesis protein NifE | 245 |  |  |
| NifE-like | Thermodesulforhabdus norvegica | K02586 | nifD; nitrogenase molybdenum-iron protein alpha chain [EC:1.18.6.1] | 112 | K02587 | 23 |
| NifE | Methanobacteriales archaeon HGW-Methanobacteriales-1* | K02587 | nifE; nitrogenase molybdenum-cofactor synthesis protein NifE | 452 |  |  |
| NifE | Methanobacterium paludis* | K02587 | nifE; nitrogenase molybdenum-cofactor synthesis protein NifE | 446 |  |  |
| NifE | Methanobrevibacter curvatus* | K02587 | nifE; nitrogenase molybdenum-cofactor synthesis protein NifE | 434 |  |  |
| NifE | Methanococcus maripaludis* | K02587 | nifE; nitrogenase molybdenum-cofactor synthesis protein NifE | 442 |  |  |
| NifE | Methanothermobacter thermautotrophicus* | K02587 | nifE; nitrogenase molybdenum-cofactor synthesis protein NifE | 317 |  |  |
| NifN-like | [Methanococcus aeolicus](https://www.kegg.jp/kegg-bin/blastkoala_result_gene_list?id=88b64dbb3f07c03bcecc1982f5ed6286e37591d7&passwd=BAHVl0&type=blastkoala&code=user&target=N%5FNif%5FMethanococcus%5Faeolicus) | [K02591](https://www.kegg.jp/entry/ko:K02591) | nifK; nitrogenase molybdenum-iron protein beta chain [EC:1.18.6.1] | 172 |  |  |
| NifN-like | [Methanofervidicoccus sp. A16](https://www.kegg.jp/kegg-bin/blastkoala_result_gene_list?id=88b64dbb3f07c03bcecc1982f5ed6286e37591d7&passwd=BAHVl0&type=blastkoala&code=user&target=N%5FNif%5FMethanofervidicoccus%5Fsp%2E%5FA16) | [K02591](https://www.kegg.jp/entry/ko:K02591) | nifK; nitrogenase molybdenum-iron protein beta chain [EC:1.18.6.1] | 211 |  |  |
| NifN-like | Methanothermococcus thermolithotrophicus | [K02591](https://www.kegg.jp/entry/ko:K02591) | nifK; nitrogenase molybdenum-iron protein beta chain [EC:1.18.6.1] | 185 |  |  |
| NifN-like | Methanothermococcus okinawensis | [K02591](https://www.kegg.jp/entry/ko:K02591) | nifK; nitrogenase molybdenum-iron protein beta chain [EC:1.18.6.1] | 170 |  |  |
| NifN | [Methanobacteriales archaeon HGW-Methanobacteriales-1](https://www.kegg.jp/kegg-bin/blastkoala_result_gene_list?id=88b64dbb3f07c03bcecc1982f5ed6286e37591d7&passwd=BAHVl0&type=blastkoala&code=user&target=N%5FNif%5FMethanobacteriales%5Farchaeon%5FHGW%2DMethanobacteriales%2D1)* | [K02591](https://www.kegg.jp/entry/ko:K02591) | nifK; nitrogenase molybdenum-iron protein beta chain [EC:1.18.6.1] | 99 | [K02592](https://www.kegg.jp/entry/ko:K02592) | 52 |
| NifN | [Methanobacterium paludis](https://www.kegg.jp/kegg-bin/blastkoala_result_gene_list?id=88b64dbb3f07c03bcecc1982f5ed6286e37591d7&passwd=BAHVl0&type=blastkoala&code=user&target=N%5FNif%5FMethanobacterium%5Fpaludis)* |  |  |  |  |  |
| NifN | Methanobrevibacter curvatus* |  |  |  |  |  |
| NifN | Methanococcus maripaludis* | [K02591](https://www.kegg.jp/entry/ko:K02591) | nifK; nitrogenase molybdenum-iron protein beta chain [EC:1.18.6.1] | 147 |  |  |
| NifN | Methanothermobacter thermoautotrophicus* | K02591 | nifK; nitrogenase molybdenum-iron protein beta chain [EC:1.18.6.1] | 116 | K02592 | 36 |

*A subset of characterized NifE and NifN homologs were included as a control.

**Table S2.** Jensen-Shannon (J-S) distances between nitrogenases and maturases at putative functionally divergent sites (see main text for definition of divergent sites), calculated for sequence alignments used to build Tree-1 to Tree-4. Site numbers from aligned *Azotobacter vinelandii* nitrogenase NifD protein. Cells are colored grey if the corresponding site is not defined as functionally divergent for the given alignment.

|  | J-S distance | | | |
| --- | --- | --- | --- | --- |
| Site | Tree-1 | Tree-2 | Tree-3 | Tree-4 |
| 15 | 0.7304 | 0.7358 | 0.73 |  |
| 16 | 0.5438 | 0.5637 | 0.5442 |  |
| 17 | 0.5687 | 0.5704 | 0.5706 | 0.566 |
| 19 |  |  |  | 0.6098 |
| 20 | 0.5462 | 0.5772 | 0.5461 | 0.6957 |
| 21 | 0.5456 | 0.5775 | 0.5455 | 0.6815 |
| 22 | 0.7661 | 0.767 | 0.7659 |  |
| 23 | 0.6701 | 0.6889 | 0.6699 | 0.5829 |
| 24 |  |  |  | 0.54 |
| 25 | 0.5837 | 0.5981 | 0.5859 | 0.6326 |
| 26 | 0.6058 | 0.6209 | 0.605 | 0.615 |
| 27 |  |  |  | 0.5254 |
| 31 | 0.5658 | 0.5953 | 0.5652 |  |
| 34 | 0.5509 | 0.5666 | 0.5499 |  |
| 41 | 0.5423 | 0.582 | 0.5419 | 0.538 |
| 43 |  |  |  | 0.6596 |
| 44 |  |  |  | 0.5749 |
| 45 |  |  |  | 0.6333 |
| 46 | 0.6614 | 0.6726 | 0.6608 |  |
| 47 |  |  |  | 0.6297 |
| 48 | 0.6955 | 0.7535 | 0.6953 | 0.6234 |
| 49 | 0.6484 | 0.6464 | 0.6481 | 0.6647 |
| 50 |  |  |  | 0.5262 |
| 51 | 0.57 | 0.5928 | 0.5701 | 0.535 |
| 52 |  | 0.5784 |  | 0.5347 |
| 56 | 0.6579 | 0.6882 | 0.6574 | 0.6462 |
| 57 | 0.7084 | 0.7332 | 0.708 | 0.687 |
| 58 |  |  |  | 0.5344 |
| 65 | 0.6453 | 0.6409 | 0.6448 | 0.6448 |
| 68 | 0.5934 | 0.6489 | 0.5929 | 0.5929 |
| 69 | 0.797 | 0.796 | 0.7969 | 0.7969 |
| 72 | 0.5676 |  | 0.5674 | 0.5674 |
| 73 | 0.7624 | 0.779 | 0.7623 | 0.7623 |
| 76 | 0.659 | 0.7087 | 0.6585 | 0.6585 |
| 78 | 0.5797 | 0.6037 | 0.5791 | 0.5791 |
| 89 | 0.5643 | 0.5959 | 0.564 | 0.5626 |
| 90 | 0.6805 | 0.7182 | 0.68 | 0.6782 |
| 94 | 0.711 | 0.717 | 0.7106 | 0.5681 |
| 95 | 0.681 | 0.6776 | 0.6805 | 0.5629 |
| 97 | 0.7381 | 0.7687 | 0.7378 | 0.5995 |
| 98 | 0.7749 | 0.7797 | 0.7747 | 0.7077 |
| 99 |  | 0.5579 |  | 0.5937 |
| 102 | 0.6329 | 0.6742 | 0.6324 |  |
| 104 | 0.6927 | 0.7335 | 0.6924 |  |
| 106 | 0.6831 | 0.7373 | 0.6827 |  |
| 108 | 0.7818 | 0.782 | 0.7817 |  |
| 109 | 0.7092 | 0.7157 | 0.7088 | 0.7096 |
| 110 | 0.7558 | 0.755 | 0.7556 | 0.7391 |
| 111 | 0.6264 | 0.6874 | 0.6258 | 0.6334 |
| 112 | 0.5977 | 0.602 | 0.5974 | 0.6044 |
| 113 | 0.6909 | 0.6922 | 0.6905 | 0.6776 |
| 118 | 0.5922 | 0.6003 | 0.5916 | 0.5916 |
| 139 | 0.5519 | 0.5696 | 0.5512 | 0.5513 |
| 141 | 0.6037 | 0.631 | 0.6033 | 0.6045 |
| 142 | 0.6541 | 0.6751 | 0.6538 | 0.6535 |
| 143 | 0.7407 | 0.7854 | 0.7404 |  |
| 144 | 0.6035 | 0.631 | 0.6032 |  |
| 149 | 0.6813 | 0.7186 | 0.6811 | 0.6811 |
| 155 | 0.7239 | 0.7618 | 0.7236 | 0.7236 |
| 167 | 0.6155 | 0.637 | 0.6199 | 0.6199 |
| 182 | 0.6057 | 0.629 | 0.6051 | 0.6051 |
| 187 | 0.745 | 0.7724 | 0.7447 |  |
| 188 | 0.7239 | 0.7789 | 0.7236 |  |
| 189 | 0.7818 | 0.7956 | 0.7817 | 0.7821 |
| 190 | 0.5563 | 0.5662 | 0.5552 | 0.5552 |
| 191 | 0.7429 | 0.8003 | 0.7425 | 0.7425 |
| 192 | 0.657 | 0.6556 | 0.6563 | 0.6563 |
| 195 | 0.7191 | 0.7697 | 0.7187 | 0.7187 |
| 196 | 0.7834 | 0.7817 | 0.7833 | 0.7833 |
| 197 | 0.5393 | 0.5559 | 0.5412 | 0.5412 |
| 199 | 0.7649 | 0.7801 | 0.7647 | 0.7647 |
| 200 | 0.5421 | 0.5689 | 0.5431 | 0.5431 |
| 205 | 0.57 | 0.576 | 0.5709 | 0.573 |
| 216 | 0.6118 | 0.606 | 0.6121 |  |
| 231 | 0.5517 |  | 0.5506 | 0.5512 |
| 232 | 0.6688 | 0.6949 | 0.6725 | 0.6712 |
| 234 | 0.6597 | 0.6785 | 0.6591 | 0.6591 |
| 235 | 0.547 | 0.5947 | 0.546 | 0.546 |
| 237 | 0.545 | 0.5839 | 0.5443 | 0.5443 |
| 253 |  |  |  | 0.5439 |
| 274 | 0.6104 | 0.6484 | 0.6097 | 0.6097 |
| 276 | 0.6672 | 0.6705 | 0.6668 | 0.6668 |
| 277 | 0.7018 | 0.7167 | 0.7014 | 0.7014 |
| 279 | 0.5522 |  | 0.5524 | 0.5524 |
| 280 | 0.6949 | 0.7157 | 0.6945 | 0.6945 |
| 281 | 0.5782 | 0.5753 | 0.5774 | 0.5774 |
| 282 | 0.5504 |  | 0.5506 | 0.5506 |
| 285 | 0.553 | 0.5742 | 0.5547 | 0.5547 |
| 294 | 0.5653 | 0.6207 | 0.5645 | 0.5645 |
| 298 | 0.6738 | 0.7358 | 0.6735 | 0.6735 |
| 302 |  | 0.5632 |  |  |
| 304 | 0.6519 | 0.6682 | 0.6519 | 0.6519 |
| 327 | 0.5514 | 0.5633 | 0.5507 | 0.5219 |
| 352 | 0.5876 | 0.6096 | 0.5909 | 0.5909 |
| 358 | 0.7199 | 0.7429 | 0.7196 | 0.7453 |
| 359 | 0.5804 | 0.6319 | 0.5801 | 0.5801 |
| 361 | 0.6183 | 0.6353 | 0.6189 | 0.6189 |
| 362 | 0.7864 | 0.7915 | 0.7863 | 0.7863 |
| 363 | 0.6387 | 0.6433 | 0.6385 | 0.6385 |
| 365 | 0.5913 | 0.6028 | 0.5905 | 0.562 |
| 379 | 0.5799 | 0.5916 | 0.58 | 0.58 |
| 380 | 0.6156 | 0.6136 | 0.6206 | 0.6206 |
| 381 | 0.7379 | 0.7954 | 0.7376 | 0.7376 |
| 383 | 0.7892 | 0.7879 | 0.7891 | 0.7891 |
| 385 |  |  |  | 0.5347 |
| 390 | 0.7294 | 0.7346 | 0.7291 | 0.6225 |
| 391 | 0.7297 | 0.7407 | 0.7293 |  |
| 393 | 0.6666 | 0.6826 | 0.666 | 0.6329 |
| 403 |  |  |  | 0.6413 |
| 404 | 0.5889 | 0.6045 | 0.5882 |  |
| 405 |  |  |  | 0.7308 |
| 406 | 0.6473 | 0.6811 | 0.6483 | 0.7155 |
| 407 | 0.571 | 0.5914 | 0.5713 | 0.5764 |
| 415 | 0.5565 | 0.5782 | 0.5558 | 0.5547 |
| 416 | 0.635 | 0.6562 | 0.635 | 0.6327 |
| 418 | 0.6885 | 0.7324 | 0.688 | 0.688 |
| 421 | 0.559 | 0.587 | 0.5584 | 0.5584 |
| 422 | 0.677 | 0.7234 | 0.6766 | 0.6766 |
| 423 | 0.5441 |  | 0.5445 | 0.5445 |
| 425 | 0.6596 | 0.6761 | 0.6592 | 0.6592 |
| 428 | 0.7035 | 0.7436 | 0.7032 | 0.684 |
| 431 |  |  |  | 0.5444 |
| 432 | 0.7118 | 0.7586 | 0.7116 | 0.7108 |
| 439 | 0.6033 | 0.6253 | 0.6035 | 0.6045 |
| 440 | 0.7852 | 0.7873 | 0.7851 | 0.7759 |
| 441 | 0.6988 | 0.7037 | 0.7004 | 0.7521 |
| 442 | 0.7998 | 0.8001 | 0.7998 |  |
| 443 | 0.7556 | 0.7598 | 0.7554 | 0.6971 |
| 444 | 0.7883 | 0.7874 | 0.7883 | 0.7927 |
| 445 | 0.7894 | 0.7934 | 0.7893 | 0.6103 |
| 446 | 0.5442 | 0.5761 | 0.5441 | 0.7547 |
| 447 | 0.6367 | 0.7113 | 0.6365 | 0.6679 |
| 448 | 0.7598 | 0.7767 | 0.7597 | 0.7597 |
| 456 |  | 0.6125 |  | 0.5321 |
| 462 | 0.5893 | 0.6013 | 0.589 | 0.589 |
| 477 |  | 0.5634 |  |  |

**Table S3.** Ancestral D-scores for putative functionally divergent sites between nitrogenases and maturases, calculated for sequence alignments used to build Tree-1 to Tree-4. Site numbers from aligned *Azotobacter vinelandii* nitrogenase NifD protein. See main text for definitions of ancestral nodes (“LCA”: Last common ancestor). Positive values are colored red and negative values are colored blue. Cells are colored grey if the corresponding site is not defined as functionally divergent for the given alignment.

|  | Nitrogenase/maturase LCA 1 | | | | Nitrogenase/maturase LCA 2 | | | | Nitrogenase LCA | | | |
| --- | --- | --- | --- | --- | --- | --- | --- | --- | --- | --- | --- | --- |
| Site | Tree-1 | Tree-2 | Tree-3 | Tree-4 | Tree-1 | Tree-2 | Tree-3 | Tree-4 | Tree-1 | Tree-2 | Tree-3 | Tree-4 |
| 15 | -0.48 | -0.16 | -0.56 |  | -0.62 | -0.15 | -0.62 |  | -0.59 | 0.09 | -0.57 |  |
| 16 | 0.11 | 0.35 | -0.12 |  | 0.07 | 0.38 | -0.15 |  | 0.11 | 0.38 | 0.24 |  |
| 17 | -0.30 | -0.14 | -0.09 | -0.41 | -0.26 | 0.11 | -0.08 | -0.22 | 0.29 | 0.45 | 0.28 | 0.10 |
| 19 |  |  |  | -0.43 |  |  |  | -0.47 |  |  |  | 0.16 |
| 20 | -0.59 | -0.61 | -0.59 | -0.44 | -0.59 | -0.61 | -0.63 | -0.42 | -0.59 | -0.60 | -0.59 | 0.35 |
| 21 | -0.39 | -0.61 | 0.38 | -0.44 | -0.57 | -0.60 | -0.63 | -0.40 | -0.60 | -0.50 | -0.59 | 0.07 |
| 22 | -0.67 | -0.70 | -0.67 |  | -0.67 | -0.67 | -0.72 |  | -0.69 | 0.07 | -0.68 |  |
| 23 | -0.32 | -0.32 | -0.33 | -0.18 | -0.31 | -0.31 | -0.33 | -0.11 | -0.32 | 0.44 | -0.32 | 0.38 |
| 24 |  |  |  | -0.28 |  |  |  | -0.30 |  |  |  | 0.18 |
| 25 | -0.38 | -0.37 | -0.03 | -0.24 | -0.44 | -0.35 | -0.26 | -0.28 | -0.28 | -0.20 | -0.26 | 0.01 |
| 26 | -0.31 | -0.10 | -0.24 | -0.24 | -0.08 | 0.16 | -0.04 | -0.27 | 0.16 | 0.25 | 0.15 | 0.21 |
| 27 |  |  |  | -0.25 |  |  |  | -0.27 |  |  |  | 0.07 |
| 31 | 0.01 | 0.29 | -0.01 |  | 0.27 | 0.45 | 0.24 |  | 0.45 | 0.45 | 0.45 |  |
| 34 | 0.02 | 0.21 | -0.01 |  | 0.17 | 0.45 | 0.20 |  | 0.41 | 0.43 | 0.40 |  |
| 41 | -0.39 | 0.30 | -0.32 | -0.61 | 0.24 | 0.21 | 0.23 | -0.61 | 0.22 | 0.21 | 0.22 | -0.65 |
| 43 |  |  |  | -0.48 |  |  |  | -0.48 |  |  |  | 0.09 |
| 44 |  |  |  | -0.44 |  |  |  | -0.44 |  |  |  | 0.18 |
| 45 |  |  |  | -0.43 |  |  |  | -0.43 |  |  |  | 0.15 |
| 46 | -0.47 | 0.05 | -0.44 |  | 0.03 | 0.11 | 0.24 |  | 0.24 | 0.13 | 0.18 |  |
| 47 |  |  |  | -0.21 |  |  |  | 0.38 |  |  |  | 0.58 |
| 48 | -0.34 | -0.44 | -0.03 | -0.09 | -0.41 | -0.62 | -0.36 | -0.39 | -0.57 | -0.56 | -0.57 | -0.36 |
| 49 | -0.02 | 0.12 | -0.03 | -0.08 | 0.03 | 0.37 | -0.05 | 0.12 | 0.44 | 0.43 | 0.41 | 0.45 |
| 50 |  |  |  | -0.12 |  |  |  | -0.15 |  |  |  | 0.22 |
| 51 | 0.16 | 0.11 | -0.27 | -0.08 | 0.13 | 0.20 | -0.40 | 0.02 | 0.20 | 0.24 | -0.21 | 0.15 |
| 52 |  | 0.42 |  | -0.07 |  | 0.45 |  | 0.39 |  | 0.56 |  | 0.36 |
| 56 | 0.30 | 0.38 | 0.34 | -0.19 | 0.28 | 0.23 | 0.29 | -0.01 | 0.23 | 0.30 | 0.25 | -0.01 |
| 57 | -0.02 | 0.03 | -0.03 | 0.44 | -0.04 | 0.06 | 0.09 | 0.24 | 0.20 | 0.31 | 0.21 | 0.22 |
| 58 |  |  |  | 0.39 |  |  |  | 0.36 |  |  |  | 0.34 |
| 65 | 0.40 | 0.42 | 0.41 | 0.16 | 0.39 | 0.42 | 0.28 | 0.22 | 0.39 | 0.45 | 0.32 | 0.44 |
| 68 | -0.02 | -0.05 | 0.00 | -0.05 | -0.06 | 0.16 | -0.28 | -0.28 | -0.28 | 0.68 | -0.28 | -0.28 |
| 69 | -0.69 | -0.71 | -0.69 | -0.69 | -0.69 | -0.71 | -0.69 | -0.69 | -0.69 | 0.43 | -0.69 | 0.44 |
| 72 | -0.19 | 0.03 | 0.00 | -0.49 | -0.49 | 0.10 | -0.62 | -0.59 | -0.59 | 0.50 | -0.59 | -0.60 |
| 73 | -0.02 | 0.43 | 0.00 | -0.03 | -0.02 | 0.38 | 0.19 | 0.00 | 0.02 | 0.37 | 0.35 | 0.51 |
| 76 | 0.40 | -0.38 | 0.41 | 0.45 | 0.40 | -0.45 | 0.55 | 0.36 | 0.36 | 0.10 | 0.36 | 0.35 |
| 78 | -0.44 | -0.33 | -0.02 | -0.37 | -0.45 | -0.34 | -0.36 | -0.48 | -0.48 | -0.22 | -0.40 | -0.10 |
| 89 | -0.37 | 0.00 | -0.36 | -0.37 | -0.36 | 0.00 | -0.40 | -0.35 | -0.35 | 0.14 | -0.35 | -0.36 |
| 90 | 0.21 | 0.11 | 0.05 | 0.20 | 0.20 | 0.06 | 0.24 | 0.19 | 0.20 | 0.08 | 0.20 | 0.19 |
| 94 | -0.01 | -0.17 | 0.09 | 0.03 | -0.01 | -0.07 | 0.14 | -0.21 | -0.02 | 0.38 | 0.02 | -0.28 |
| 95 | -0.21 | 0.70 | 0.04 | 0.29 | -0.23 | 0.72 | -0.12 | 0.29 | -0.15 | 0.71 | -0.14 | 0.29 |
| 97 | -0.03 | -0.06 | 0.00 | 0.17 | 0.00 | 0.06 | 0.24 | 0.28 | 0.25 | 0.37 | 0.13 | 0.42 |
| 98 | -0.24 | 0.04 | -0.01 | 0.25 | -0.29 | -0.03 | -0.52 | 0.15 | 0.01 | -0.01 | -0.18 | 0.28 |
| 99 |  |  |  | -0.33 |  |  |  | -0.29 |  |  |  | -0.23 |
| 102 | 0.15 | -0.21 | 0.12 |  | -0.19 | -0.56 | -0.11 |  | -0.43 | -0.40 | -0.51 |  |
| 104 | -0.61 | -0.09 | -0.57 |  | -0.66 | -0.07 | -0.69 |  | -0.57 | 0.36 | -0.57 |  |
| 106 | 0.34 | -0.51 | 0.35 |  | -0.54 | -0.32 | -0.52 |  | -0.17 | -0.31 | -0.42 |  |
| 108 | 0.00 | 0.04 | 0.08 |  | -0.73 | 0.15 | -0.76 |  | -0.74 | 0.39 | -0.71 |  |
| 109 | -0.23 | -0.34 | -0.01 | -0.04 | -0.29 | -0.45 | -0.13 | -0.05 | -0.20 | -0.12 | -0.17 | 0.07 |
| 110 | -0.22 | -0.37 | 0.01 | -0.06 | -0.50 | -0.32 | -0.46 | -0.06 | -0.30 | 0.20 | -0.30 | -0.06 |
| 111 | -0.19 | -0.23 | 0.00 | -0.12 | -0.53 | -0.22 | -0.17 | -0.16 | -0.49 | -0.07 | -0.43 | -0.03 |
| 112 | -0.10 | -0.12 | -0.02 | -0.07 | -0.10 | 0.01 | -0.14 | -0.09 | 0.20 | 0.34 | 0.09 | 0.36 |
| 113 | 0.04 | 0.04 | 0.00 | 0.02 | 0.06 | 0.09 | 0.22 | 0.06 | 0.17 | 0.10 | 0.14 | 0.18 |
| 118 | -0.42 | -0.30 | -0.47 | -0.44 | -0.44 | -0.29 | -0.54 | -0.42 | -0.44 | -0.09 | -0.42 | 0.24 |
| 139 | -0.27 | -0.02 | -0.05 | -0.04 | -0.25 | 0.00 | -0.13 | -0.17 | -0.19 | 0.23 | -0.21 | 0.16 |
| 141 | -0.22 | -0.26 | -0.14 | -0.32 | -0.24 | -0.25 | -0.21 | -0.20 | -0.09 | -0.32 | 0.06 | -0.21 |
| 142 | -0.26 | -0.32 | 0.00 | -0.27 | -0.27 | -0.26 | 0.07 | 0.26 | 0.45 | 0.42 | 0.33 | 0.50 |
| 143 | -0.63 | -0.71 | -0.62 |  | -0.63 | -0.67 | -0.75 |  | 0.30 | 0.31 | -0.37 |  |
| 144 | -0.42 | -0.39 | -0.09 |  | -0.44 | -0.19 | -0.34 |  | -0.21 | -0.11 | -0.11 |  |
| 149 | -0.60 | -0.63 | -0.59 | -0.62 | -0.59 | -0.61 | -0.61 | -0.58 | -0.58 | -0.52 | -0.58 | -0.59 |
| 155 | 0.16 | 0.08 | 0.02 | 0.08 | 0.15 | 0.05 | 0.06 | 0.08 | 0.24 | 0.10 | 0.05 | 0.14 |
| 167 | -0.64 | -0.60 | -0.13 | -0.61 | -0.62 | -0.54 | -0.72 | -0.48 | -0.49 | -0.16 | -0.50 | -0.10 |
| 182 | 0.23 | 0.07 | 0.18 | 0.11 | 0.15 | 0.07 | -0.01 | 0.07 | 0.07 | 0.07 | 0.07 | 0.07 |
| 187 | -0.57 | -0.65 | -0.57 |  | -0.58 | -0.66 | -0.40 |  | 0.37 | 0.35 | 0.27 |  |
| 188 | -0.22 | -0.13 | -0.01 |  | -0.19 | -0.01 | 0.33 |  | 0.71 | 0.72 | 0.71 |  |
| 189 | -0.61 | -0.68 | -0.02 | -0.65 | -0.61 | -0.69 | -0.02 | 0.07 | 0.32 | 0.25 | 0.33 | 0.48 |
| 190 | 0.38 | 0.38 | -0.15 | 0.44 | 0.40 | 0.41 | 0.24 | 0.65 | 0.65 | 0.65 | 0.65 | 0.65 |
| 191 | -0.02 | -0.02 | 0.00 | -0.02 | -0.02 | -0.07 | -0.03 | 0.00 | 0.00 | 0.00 | -0.01 | 0.00 |
| 192 | 0.09 | 0.07 | -0.02 | 0.08 | 0.19 | 0.57 | 0.37 | 0.61 | 0.72 | 0.71 | 0.71 | 0.71 |
| 195 | -0.13 | -0.07 | 0.00 | -0.14 | -0.09 | 0.68 | 0.60 | 0.73 | 0.73 | 0.73 | 0.72 | 0.72 |
| 196 | -0.06 | -0.03 | -0.04 | -0.06 | -0.07 | 0.29 | -0.10 | -0.14 | -0.11 | 0.75 | -0.27 | 0.75 |
| 197 | -0.27 | -0.13 | -0.18 | -0.05 | -0.17 | -0.02 | 0.06 | 0.25 | 0.24 | 0.31 | 0.25 | 0.25 |
| 199 | -0.04 | -0.09 | 0.00 | -0.05 | -0.07 | -0.06 | -0.06 | -0.07 | -0.04 | 0.75 | 0.01 | 0.73 |
| 200 | -0.15 | -0.15 | -0.23 | -0.13 | -0.13 | 0.17 | -0.10 | 0.00 | 0.03 | 0.24 | 0.04 | 0.16 |
| 205 | 0.14 | 0.14 | 0.12 | -0.06 | 0.14 | 0.04 | 0.07 | 0.01 | 0.01 | 0.07 | 0.02 | 0.02 |
| 216 | -0.43 | -0.30 | -0.42 |  | -0.47 | -0.31 | -0.53 |  | -0.46 | 0.06 | -0.44 |  |
| 231 | -0.26 |  | -0.11 | -0.28 | -0.16 |  | -0.11 | -0.09 | -0.01 |  | -0.10 | 0.49 |
| 232 | -0.14 | 0.05 | -0.18 | -0.03 | -0.20 | 0.01 | -0.51 | -0.48 | -0.42 | 0.08 | -0.50 | -0.11 |
| 234 | 0.70 | 0.70 | 0.00 | 0.70 | 0.70 | 0.70 | 0.71 | 0.70 | 0.70 | 0.70 | 0.70 | 0.70 |
| 235 | -0.21 | -0.25 | 0.00 | -0.22 | -0.21 | -0.16 | -0.28 | -0.12 | -0.12 | -0.17 | -0.12 | -0.12 |
| 237 | -0.09 | -0.09 | 0.10 | -0.08 | -0.08 | -0.06 | 0.03 | 0.14 | 0.14 | -0.05 | 0.16 | 0.13 |
| 253 |  |  |  | -0.19 |  |  |  | 0.17 |  |  |  | 0.17 |
| 274 | -0.02 | 0.02 | 0.00 | -0.02 | -0.02 | 0.22 | 0.19 | 0.18 | 0.17 | 0.28 | 0.17 | 0.17 |
| 276 | 0.03 | 0.06 | 0.00 | 0.02 | 0.01 | 0.04 | 0.00 | 0.01 | 0.03 | 0.08 | 0.01 | 0.08 |
| 277 | -0.10 | -0.09 | 0.00 | -0.07 | -0.12 | 0.06 | -0.18 | 0.23 | 0.11 | 0.60 | -0.12 | 0.66 |
| 279 | 0.10 |  | -0.13 | -0.05 | 0.09 |  | -0.18 | -0.03 | 0.11 |  | -0.09 | 0.09 |
| 280 | 0.00 | 0.02 | 0.01 | -0.07 | -0.03 | 0.14 | -0.17 | 0.01 | 0.02 | 0.18 | -0.06 | 0.02 |
| 281 | -0.26 | -0.31 | -0.17 | -0.33 | -0.19 | 0.65 | -0.21 | 0.66 | 0.69 | 0.68 | 0.69 | 0.68 |
| 282 | -0.17 |  | 0.01 | -0.20 | -0.27 |  | -0.30 | -0.33 | -0.23 |  | -0.31 | 0.33 |
| 285 | -0.29 | -0.36 | -0.05 | -0.10 | -0.38 | -0.36 | -0.47 | -0.12 | -0.09 | -0.11 | -0.34 | 0.15 |
| 294 | -0.41 | -0.50 | -0.01 | -0.40 | -0.40 | -0.25 | -0.41 | -0.18 | -0.17 | -0.21 | -0.17 | -0.18 |
| 298 | -0.01 | -0.12 | -0.01 | -0.12 | -0.11 | -0.10 | -0.16 | -0.55 | -0.58 | 0.50 | -0.56 | -0.58 |
| 302 |  | -0.46 |  |  |  | -0.48 |  |  |  | -0.41 |  |  |
| 304 | -0.05 | -0.19 | 0.03 | 0.02 | -0.07 | -0.38 | 0.10 | -0.09 | -0.09 | -0.31 | -0.10 | -0.09 |
| 327 | 0.00 | -0.12 | 0.00 | -0.27 | -0.03 | 0.06 | 0.44 | 0.43 | 0.63 | 0.64 | 0.50 | 0.54 |
| 352 | -0.22 | -0.07 | 0.26 | -0.24 | -0.23 | 0.00 | -0.17 | -0.20 | -0.20 | 0.40 | -0.16 | 0.28 |
| 358 | -0.15 | -0.18 | -0.03 | 0.14 | -0.15 | -0.17 | 0.17 | 0.27 | 0.21 | -0.02 | 0.23 | 0.28 |
| 359 | 0.01 | -0.10 | 0.00 | 0.40 | 0.04 | -0.27 | 0.59 | 0.60 | 0.65 | -0.07 | 0.52 | 0.50 |
| 361 | -0.24 | -0.32 | -0.13 | -0.28 | -0.28 | -0.09 | -0.29 | -0.13 | -0.05 | -0.09 | -0.05 | -0.05 |
| 362 | -0.32 | -0.40 | 0.00 | -0.39 | -0.48 | -0.46 | -0.63 | -0.64 | -0.63 | 0.75 | -0.59 | 0.75 |
| 363 | -0.37 | -0.39 | -0.28 | -0.40 | -0.38 | -0.34 | -0.38 | -0.20 | -0.24 | -0.33 | -0.15 | 0.14 |
| 365 | -0.15 | -0.18 | -0.09 | -0.18 | -0.18 | -0.23 | -0.21 | -0.15 | -0.18 | -0.21 | -0.16 | -0.13 |
| 379 | -0.17 | -0.24 | -0.17 | -0.21 | -0.18 | -0.25 | -0.28 | -0.23 | -0.19 | -0.28 | -0.18 | -0.18 |
| 380 | 0.17 | 0.21 | 0.00 | 0.04 | 0.23 | 0.39 | 0.24 | 0.35 | 0.34 | 0.35 | 0.35 | 0.23 |
| 381 | -0.03 | 0.02 | 0.00 | -0.01 | -0.03 | 0.09 | 0.47 | 0.74 | 0.74 | 0.74 | 0.73 | 0.73 |
| 383 | -0.27 | 0.47 | 0.25 | -0.17 | -0.28 | 0.53 | -0.03 | -0.15 | -0.12 | 0.74 | 0.08 | 0.77 |
| 385 |  |  |  | 0.00 |  |  |  | 0.04 |  |  |  | 0.30 |
| 390 | 0.09 | 0.22 | 0.02 | -0.03 | 0.18 | 0.28 | 0.11 | 0.01 | -0.68 | 0.24 | -0.68 | -0.02 |
| 391 | -0.64 | -0.02 | -0.62 |  | -0.67 | -0.02 | -0.73 |  | -0.64 | 0.37 | -0.64 |  |
| 393 | -0.36 | 0.00 | -0.32 | -0.30 | -0.24 | 0.17 | 0.10 | -0.22 | 0.17 | 0.27 | 0.18 | 0.14 |
| 403 |  |  |  | 0.60 |  |  |  | 0.55 |  |  |  | 0.04 |
| 404 | -0.19 | -0.19 | -0.02 |  | -0.19 | -0.11 | -0.31 |  | -0.11 | 0.00 | -0.11 |  |
| 405 |  |  |  | -0.65 |  |  |  | -0.66 |  |  |  | 0.29 |
| 406 | 0.01 | -0.01 | 0.02 | -0.64 | -0.03 | -0.01 | -0.03 | -0.65 | -0.03 | 0.02 | -0.03 | 0.08 |
| 407 | 0.38 | 0.37 | 0.00 | 0.45 | 0.40 | 0.35 | 0.01 | 0.03 | 0.21 | 0.38 | 0.04 | 0.08 |
| 415 | -0.11 | -0.12 | -0.01 | -0.12 | -0.10 | -0.05 | 0.13 | 0.05 | 0.06 | -0.04 | 0.03 | 0.05 |
| 416 | -0.02 | -0.11 | 0.41 | 0.09 | -0.04 | 0.08 | 0.11 | 0.03 | 0.03 | 0.46 | 0.03 | 0.04 |
| 418 | 0.68 | 0.71 | 0.66 | 0.69 | 0.69 | 0.67 | 0.69 | 0.66 | 0.66 | 0.66 | 0.66 | 0.66 |
| 421 | -0.10 | 0.04 | 0.26 | 0.01 | -0.27 | -0.01 | 0.03 | 0.03 | 0.04 | 0.15 | 0.04 | 0.05 |
| 422 | -0.52 | -0.40 | -0.42 | -0.56 | -0.57 | -0.51 | -0.66 | -0.40 | -0.44 | -0.31 | -0.35 | -0.47 |
| 423 | -0.30 |  | 0.11 | -0.29 | -0.30 |  | -0.38 | -0.32 | -0.32 |  | -0.05 | 0.18 |
| 425 | -0.08 | -0.02 | 0.03 | -0.04 | -0.08 | 0.10 | 0.08 | -0.04 | -0.04 | 0.36 | 0.09 | 0.29 |
| 428 | 0.40 | 0.39 | 0.00 | 0.40 | 0.40 | 0.40 | 0.21 | 0.39 | 0.39 | 0.40 | 0.39 | 0.39 |
| 431 |  |  |  | -0.52 |  |  |  | -0.43 |  |  |  | -0.49 |
| 432 | 0.17 | 0.07 | 0.00 | 0.10 | -0.04 | 0.01 | -0.03 | 0.00 | 0.00 | 0.08 | 0.00 | -0.01 |
| 439 | -0.14 | -0.31 | -0.01 | -0.12 | -0.15 | -0.43 | -0.18 | -0.11 | -0.11 | -0.31 | -0.15 | -0.11 |
| 440 | -0.46 | -0.57 | -0.01 | 0.03 | -0.48 | -0.56 | -0.46 | 0.03 | -0.04 | -0.05 | -0.39 | 0.08 |
| 441 | 0.03 | 0.12 | 0.00 | 0.02 | 0.11 | 0.02 | 0.01 | -0.01 | -0.01 | 0.03 | -0.05 | 0.15 |
| 442 | -0.61 | -0.46 | 0.00 |  | -0.56 | -0.39 | -0.40 |  | 0.27 | 0.73 | -0.29 |  |
| 443 | 0.22 | 0.58 | 0.00 | 0.18 | 0.23 | 0.41 | 0.16 | 0.20 | 0.15 | 0.39 | 0.07 | 0.38 |
| 444 | 0.02 | 0.36 | 0.00 | 0.25 | 0.01 | 0.38 | -0.08 | 0.07 | -0.01 | 0.38 | -0.02 | 0.39 |
| 445 | -0.34 | -0.19 | -0.03 | 0.04 | -0.35 | -0.18 | -0.16 | 0.13 | -0.08 | 0.28 | -0.15 | 0.15 |
| 446 | -0.58 | -0.63 | -0.57 | -0.29 | -0.58 | -0.63 | -0.57 | -0.33 | -0.59 | -0.63 | -0.57 | -0.40 |
| 447 | 0.08 | 0.12 | 0.00 | 0.10 | 0.12 | 0.14 | 0.11 | 0.12 | 0.16 | 0.15 | 0.05 | 0.07 |
| 448 | -0.14 | -0.20 | -0.02 | -0.09 | -0.14 | -0.08 | -0.08 | 0.00 | 0.04 | 0.75 | 0.00 | 0.70 |
| 456 |  | 0.00 |  | 0.05 |  | -0.01 |  | 0.16 |  | 0.16 |  | 0.15 |
| 462 | -0.10 | -0.07 | -0.01 | -0.08 | -0.12 | 0.15 | -0.01 | 0.43 | 0.39 | 0.44 | 0.43 | 0.48 |
| 477 |  | -0.37 |  |  |  | -0.27 |  |  |  | -0.20 |  |  |

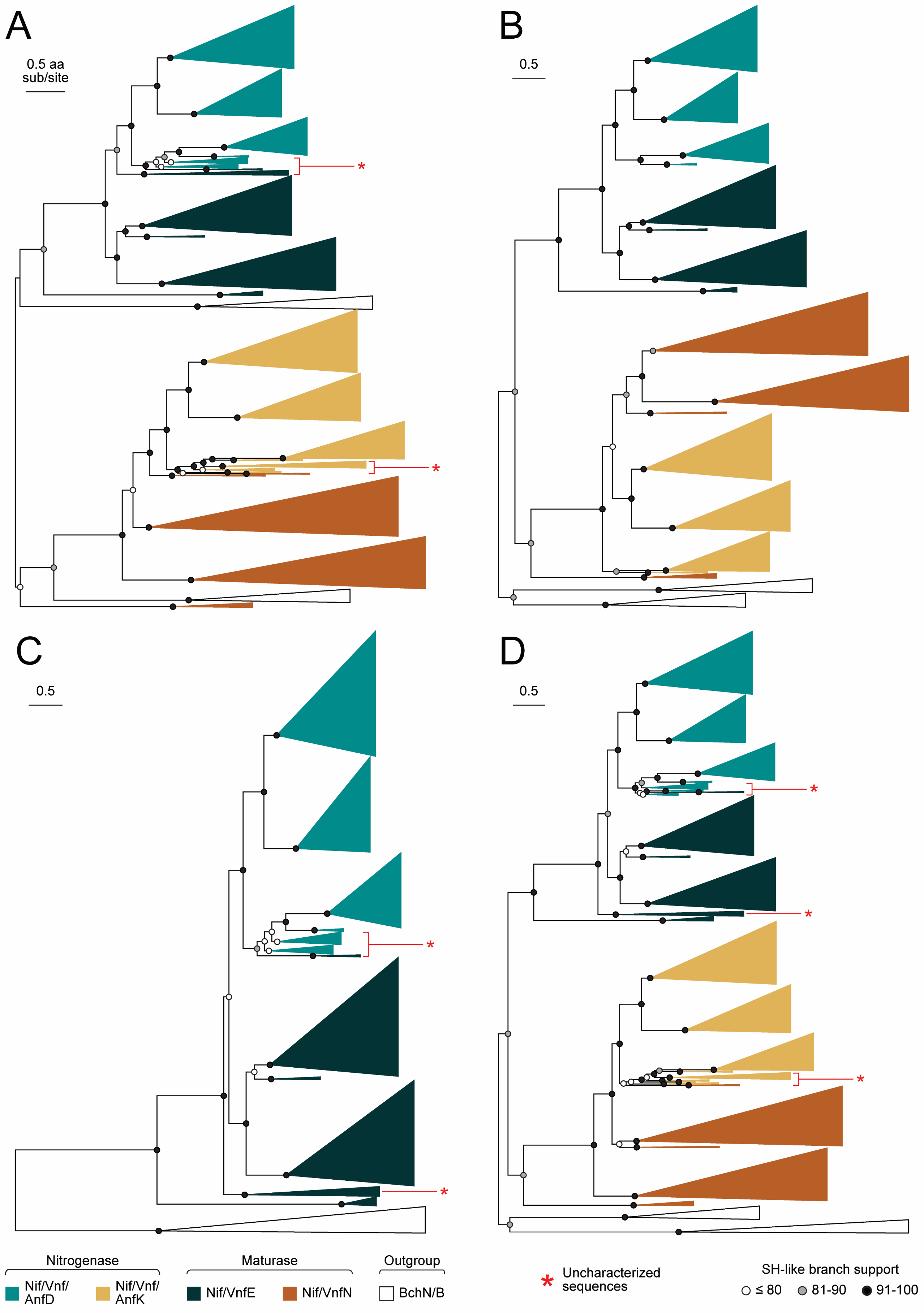

**Fig. S1.** Alternate maximum likelihood phylogenies built from nitrogenase and maturase protein sequences and incorporated into ancestral sequence inference (see main text for tree reconstruction methods): (*A*) Tree-1, (*B*) Tree-2, (*C*) Tree-3, and (*D*) Tree-4. Outgroup clade widths are not to scale.

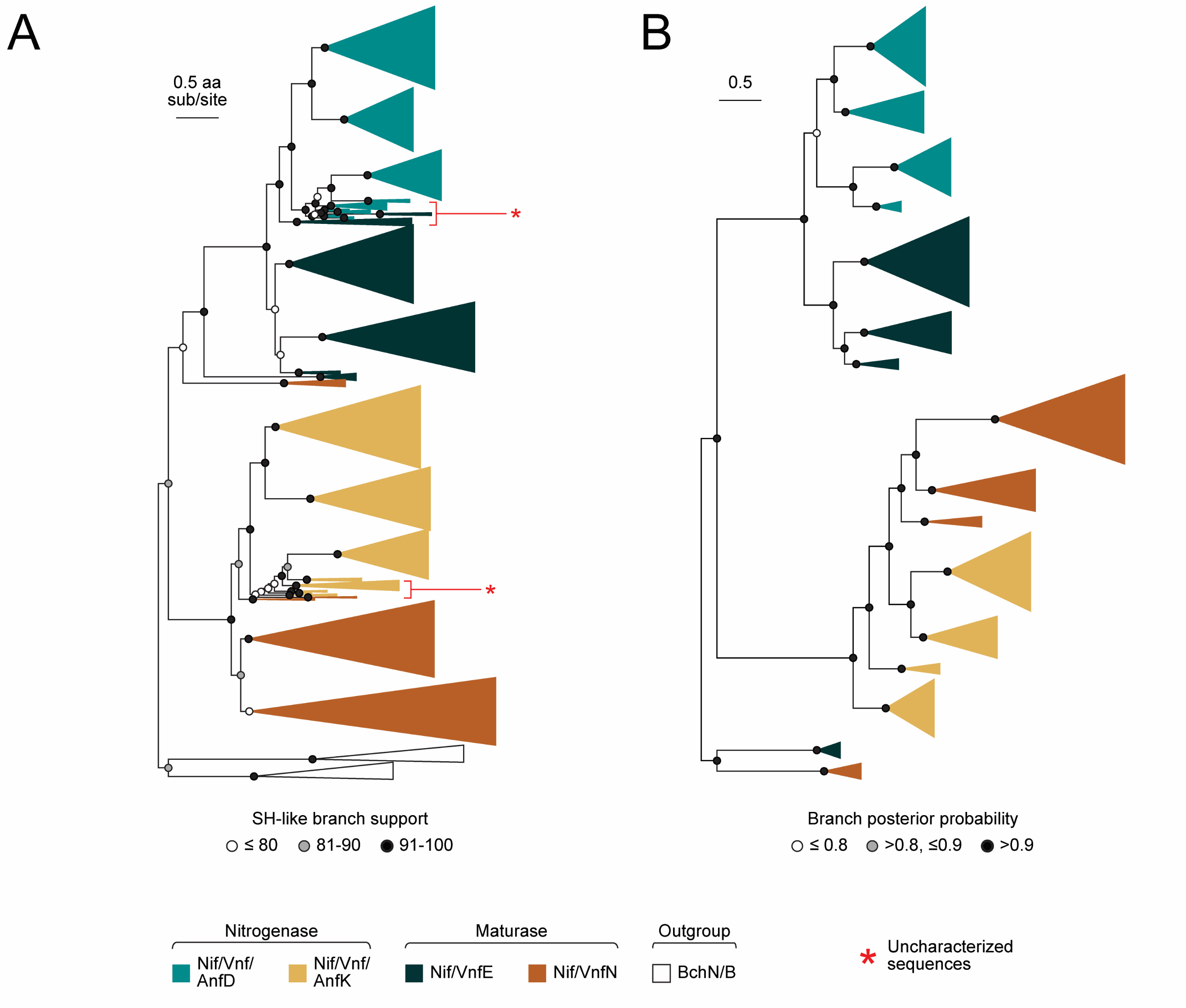

**Fig. S2.** Nitrogenase and maturase protein sequence phylogenies built (*A*) to test the effect of alignment trimming and (B) to replicate the Bayesian phylogenetic analysis by Boyd et al. (2011) with a smaller dataset lacking uncharacterized sequences (see main text for tree reconstruction methods). Outgroup clade widths are not to scale.
